# Astrocytes regulate spatial memory in a sex-specific manner

**DOI:** 10.1101/2022.11.03.511881

**Authors:** Samantha M. Meadows, Fernando Palaguachi, Avital Licht-Murava, Daniel Barnett, Till S. Zimmer, Constance Zhou, Samantha R. McDonough, Adam L. Orr, Anna G. Orr

## Abstract

Cognitive processes and neurocognitive disorders are regulated by astrocytes and have prominent sex differences. However, the contribution of astrocytes to sex differences is not known. We leveraged astrocyte-targeted gene editing and chemogenetics in adult mice to reveal that astrocytic glutamate receptors and other G protein-coupled receptors (GPCRs) modulate hippocampus-dependent cognitive function in a sexually dimorphic manner. In females, spatial memory was improved by increasing metabotropic glutamate receptor 3 (mGluR3) in astrocytes or stimulating astrocytic G_i/o_-coupled signaling, whereas stimulating G_s_-coupled signaling impaired memory. However, in males, memory was improved by reducing mGluR3 or stimulating G_s_-coupled signaling, whereas stimulating G_i/o_-coupled signaling impaired memory. Thus, memory requires a sex-specific balance of astrocytic G_s_-coupled and G_i/o_-coupled receptor activities, and disease-associated alterations or therapeutic targeting of these pathways may cause opposing sex-dependent effects on cognitive function.

**Summary:** Glia cause sex-specific changes in cognition

## Main Text

Astrocytes are abundant non-neuronal cells in the central nervous system that enable and regulate various neural functions, including cognition (*1, 2*). Astrocytic G protein-coupled receptors (GPCRs) are context-dependent modulators of memory and other neurocognitive outcomes in physiological and pathological conditions (*3-12*). Substantial evidence demonstrates sex differences in learning and memory (*13-16*), neurocognitive disorders (*17*), and aging-related cognitive decline (*15*), but it is not known if astrocytic receptor signaling contributes to these sex differences. Previous studies on the roles of astrocytic GPCRs in cognition and behavior have included only male subjects (*3, 5-12, 18, 19*) or were underpowered to assess sex differences (*4*). Most striking is that these limitations (male-only or underpowering) are common in biological research, with several groups highlighting the necessity to consider biological sex in forthcoming studies (*17, 20-22*).

We explored whether hippocampal astrocytes regulate learning and memory differently in males and females by using *in vivo* CRISPR/Cas9-based gene editing, chemogenetics, and other complementary approaches. Our findings reveal that astrocytic glutamate receptors and related types of GPCRs cause opposing, sexually dimorphic effects on memory. These results highlight novel context-dependent roles of astrocytic signaling in behavior and reveal that biological sex strongly influences astrocytic control of cognitive function.

Metabotropic glutamate receptor 3 (mGluR3) is a G_i/o_-coupled GPCR that is highly enriched in hippocampal and cortical astrocytes relative to other neural cell types (*23-25*). Alterations in mGluR3 are observed in diverse neurological diseases, including Alzheimer’s disease (*26-28*), neuropsychiatric disorders (*29, 30*), and glioblastoma (*31*). Notably, astrocytic mGluR3 levels are reduced in aging and neurocognitive disorders (*26, 32*) and single nucleotide polymorphisms in the gene encoding mGluR3 (*Grm3*) are associated with cognitive dysfunction and neuropsychiatric disorders, including schizophrenia and bipolar disorder (*29, 30, 33*). Despite strong associations with neurocognitive disorders, the exact roles of astrocytic mGluR3 in cognitive processes are not known. Therefore, we tested whether changes in astrocytic mGluR3 expression levels influence hippocampus-dependent cognitive functions, including learning and memory.

To reduce astrocytic mGluR3 expression using CRISPR/Cas9-based gene editing, we generated doubly transgenic *Aldh1l1*-CreERT2:LSL-Cas9-eGFP mice (i.e., *Aldh1l1*-Cas9) that enabled astrocyte-selective, tamoxifen (TAM)-inducible, and Cre-dependent Cas9-eGFP expression in approximately 20% of hippocampal astrocytes in adult males and females (**Fig. S1**). To selectively disrupt the gene encoding mGluR3 (*Grm3*), the mice received intrahippocampal injections of PHP.eB AAV vectors (*34, 35*) encoding custom *Grm3*-targeting sgRNAs one week prior to TAM administration. This approach resulted in a partial loss of mGluR3 expression due to the mosaic pattern of Cas9-eGFP induction in hippocampal astrocytes (**Figs. 1A**–**1B, S1, S2A**). In doubly transgenic male and female *Aldh1l1*-Cas9 mice, mGluR3-immunoreactive areas and intensities in the hippocampus were reduced by 45% and bulk *Grm3* mRNA levels were reduced by 25% (**Figs. 1C**–**1D, S2B**). At 5–7 months of age, baseline mGluR3 levels (mRNA and protein) were similar between sexes (**Fig. S2C**). There were no detectable sgRNA-induced changes of other types of mGluRs in the dentate gyrus of males and females (**Fig. S2D**–**S2E**), indicating that the knockdown was highly selective to *Grm3* and there were no compensatory alterations in other closely related metabotropic glutamate receptors, as has been reported in constitutive global mGluR3 knockout mice (*36*).

**Fig. 1.**
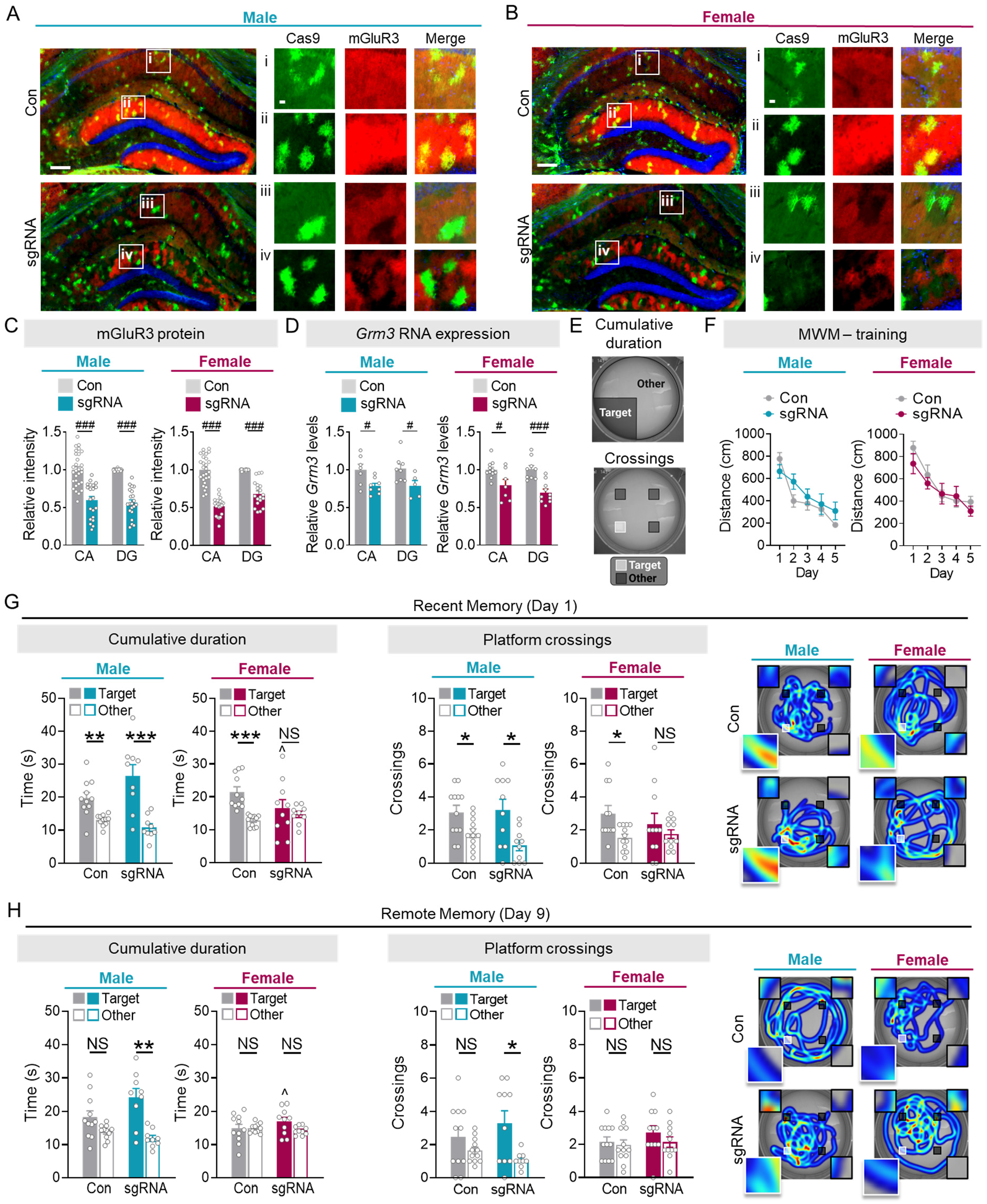
Reductions in astrocytic mGluR3 expression cause opposing sex-specific effects on memory. Representative images of co-immunolabeling for Cas9-eGFP (green) and mGluR3 (red) in hippocampal sections from tamoxifen-treated *Aldh1l1*-Cas9 male (**A**) and female (**B**) mice injected with saline (Con) or AAVs encoding mGluR3-targeting sgRNAs. Scale bars: 400 μm, 40 μm (insets). (**C**) Quantification of mGluR3 immunofluorescence in the CA1 and dentate gyrus (DG) regions of the hippocampal formation. Data for each brain region were normalized per sex to determine the extent of mGluR3 knockdown in males versus females. See Fig. S2C for basal mGluR3 levels across hippocampal regions and sexes. Two-way ANOVA (males): *F*(1, 99) = 138.8, p < 0.001 for main effect of mGluR3 manipulation. Two-way ANOVA (females): *F*(1, 90) = 194.0, p < 0.001 for main effect of mGluR3 manipulation. (**D**) Hippocampal *Grm3* mRNA levels. Two-way ANOVA (males): *F*(1, 24) = 12.61, p = 0.0016 for main effect of mGluR3 manipulation. Two-way ANOVA (females): *F*(1, 33) = 26.52, p < 0.001 for main effect of mGluR3 manipulation. (**E**) Schematic denoting areas used for assessing target and other arena areas for cumulative duration and crossings. (**F**) Mean distance traveled per training day in the Morris water maze. (**G**) Target quadrant preference and platform crossings compared to other analogous locations during the probe trial performed one day after training. Two-way ANOVA (target duration): *F*(1, 38) = 6.296, p = 0.0165 for interaction effect. Two-way ANOVA (target crossings): *F*(1, 40) = 2.166, p = 0.1489 for interaction effect. (**H**) Target quadrant preference and platform crossings in a probe trial performed 9 days after training. Two-way ANOVA (target duration): *F*(1, 37) = 1.054, p = 0.3113 for interaction effect. Two-way ANOVA (target crossings): *F*(1, 37) = 1.054, p = 0.3113 for interaction effect. Three-way ANOVA (F) or two-way ANOVA (C–D and G–H) with Sidak’s multiple comparisons post-hoc tests (vs control/sex): #p < 0.05, ###p < 0.001; post-hoc tests (vs male target/condition): ^p < 0.05. Target preference in the MWM was determined using Student’s t-test (G–H). *p < 0.05, **p < 0.01, ***p < 0.001. NS: no significant preference for target.

To evaluate the effects of astrocytic mGluR3 knockdown on hippocampus-dependent cognitive function, we used the Morris water maze (MWM) to assess spatial learning and memory in recent (one day post-training) and remote (nine days post-training) probe trials. We coupled this approach with automated behavioral analyses, which enabled unbiased and multifaceted assessments of 1) spatial learning over time, 2) preferences for target zones (i.e., target quadrants and target platform locations) relative to other areas (**Fig. 1E**), 3) adoption of specific types of spatial search strategies, 4) accuracy of starting trajectories used to approach the target, and 5) motor abilities (*37-41*). In addition to its central roles in learning and memory, the hippocampus also facilitates novelty-processing and anxiety (*42, 43*), which we also assessed using the elevated plus maze (EPM) (*44, 45*) (**Fig. S2F**).

At 5–7 months of age, knockdown of astrocytic mGluR3 did not affect learning or swimming abilities in males or females (**Figs. 1F, S3A**–**S3B**). However, knockdown of astrocytic mGluR3 impaired recent memory specifically in females, without significantly affecting search strategies or trajectories (**Figs. 1G, S3C, S3E**). In contrast, male mice with reduced mGluR3 levels had intact recent memory and enhanced remote memory as well as improved search strategies and trajectories (**Figs. 1H, S3D, S3F**, and **Supplemental text**), demonstrating that reductions in astrocytic mGluR3 have opposing and persistent sex-dependent effects on spatial memory. In the EPM, astrocytic mGluR3 knockdown reduced novelty-related exploration in females (**Fig. S3G**), but increased novelty-related exploration in males, consistent with previous findings in male mice with global mGluR3 gene ablation (*46*). Anxiety-like behavior was not affected in males or females (**Fig. S3H**). Thus, reductions in astrocytic mGluR3 had sexually dimorphic effects on hippocampus-dependent behavior in two different test paradigms, suggesting that the divergent effects are consistent across different behavioral modalities.

To further assess if astrocytic mGluR3 modulates memory, we used astrocyte-targeted AAV vectors encoding mGluR3 to test whether enhancing astrocytic mGluR3 expression also alters memory in a sex-specific manner. We delivered AAV vectors encoding Cre-dependent mGluR3 under the control of the astrocytic promoter *hGfaABC*_*1*_*D* (*47*) into the hippocampus of *Aldh1l1*-CreERT2 mice. This stringent system enabled inducible, high-efficiency, and cell type-specific targeting of mGluR3 to hippocampal astrocytes (**Figs. 2A**–**2B, S4**–**S5, S6A** and **Supplemental text**). Intrahippocampal injections of the mGluR3-encoding AAV induced a four-fold increase in mGluR3 protein levels in males and females, with similar coverage of hippocampal areas among the sexes (**Fig. 2C**–**2D**). In primary astrocyte cultures, the AAV vector encoding mGluR3 increased the levels of mGluR3 monomers and functional dimers (**Fig. S6B**) (*48, 49*). In addition, the mGluR2/3-selective agonist LY354740 increased phosphorylated Akt levels in transduced astrocytes (**Fig. S6C**), demonstrating intact surface expression and signaling by the exogenous receptor.

**Fig. 2.**
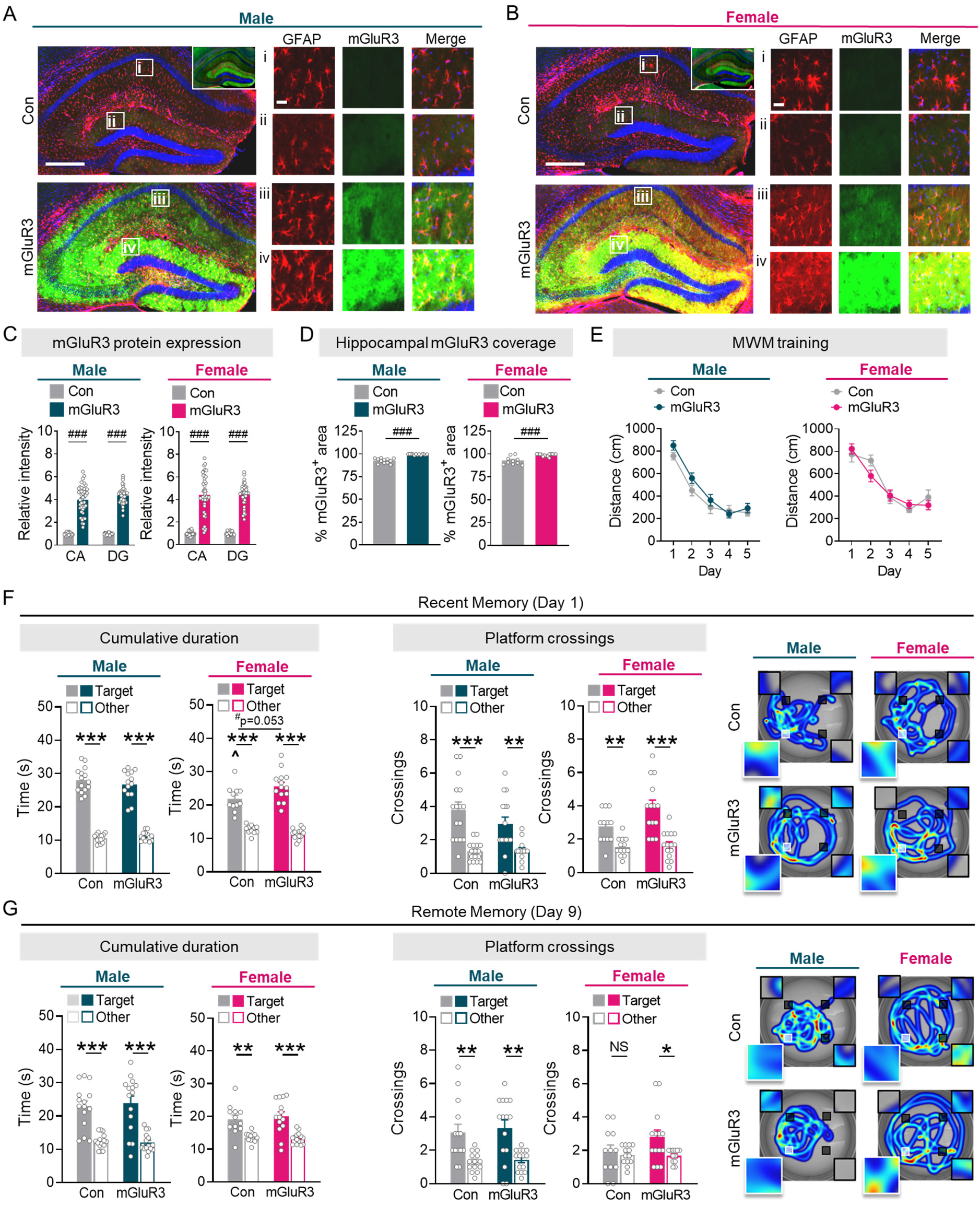
Increases in astrocytic mGluR3 expression improves memory recall in females but not males. Representative images of co-immunolabeling for GFAP (red) and mGluR3 (green) in hippocampal sections from TAM-treated *Aldh1l1*-CreERT males (**A**) and females (**B**) injected with saline (Con) or AAVs encoding mGluR3 (mGluR3). Scale bars: 400 μm; 40 μm (insets). Inlays show the same sections at higher exposure (see related data in Fig. S5). (**C**) Quantification of mGluR3 immunofluorescence in the CA1 and dentate gyrus (DG) regions of the hippocampal formation. Two-way ANOVA (males): *F*(1, 141) = 622.4, p < 0.001 for main effect of mGluR3 manipulation. Two-way ANOVA (females): *F*(1, 141) = 368.3, p < 0.001 for main effect of mGluR3 manipulation. (**D**) Quantification of the % DG area that was mGluR3-immunopositive. Two-way ANOVA: *F*(1, 45) = 86.50, p = p < 0.001 for main effect of mGluR3 manipulation. (**E**) Mean distance traveled per training day in the Morris water maze. (**F**) Target quadrant preference and platform crossings compared to other analogous locations during the probe trial performed one day after training. Two-way ANOVA (target duration): *F*(1, 50) = 4.874, p = 0.0319 for interaction effect. Two-way ANOVA (target crossings): *F*(1, 53) = 5.963, p = 0.0180 for interaction effect. (**G**) Target quadrant preference and platform crossings in a probe trial performed 9 days after training. Two-way ANOVA (target duration): *F*(1, 52) = 0.0008, p = 0.9781 for interaction effect. Two-way ANOVA (target crossings): *F*(1, 51) = 0.4293, p = 0.5153 for interaction effect. Three-way ANOVA (E) or two-way ANOVA (C–D and F–G) with Sidak’s multiple comparisons post-hoc tests (vs control/sex): ###p < 0.001; post-hoc tests (vs male target/condition): ^p < 0.05. Target preference in the MWM was determined using Student’s t-test (F–G). *p < 0.05, **p < 0.01, ***p < 0.001. NS: no significant preference for target.

Given our findings that mGluR3 reductions impaired memory in females (**Fig. 1G**), we hypothesized that mGluR3 enhancement would improve memory in females. Notably, female mice at 12–14 months of age start to develop aging-related memory loss, whereas males develop memory loss at later ages (*15*). At these ages, basal mGluR3 protein levels were already reduced in females as compared to males (**Fig. S6D**), consistent with previous reports of aging and disease-related depletions of mGluR3 in humans and mouse models of disease (*26-28, 50*). Thus, we tested whether enhancing astrocytic mGluR3 expression can improve memory in 12–14-month-old females (**Fig. S6E**). Indeed, astrocytic mGluR3 enhancement in females caused sex-specific improvements in recent and remote memory without affecting learning or swimming abilities (**Figs. 2E**–**2G, S7A**–**S7F**, and **Supplemental text**), novelty-induced exploration (**Fig. S7G**), or anxiety-linked behavior (**Fig. S7H**). Male controls maintained robust target preferences throughout testing, whereas female controls with lower basal mGluR3 expression than males had reduced target preferences and more inefficient search strategies (**Figs. 2F**–**2G, S7D**), consistent with the observed sex-specific memory impairments in females upon knockdown of astrocytic mGluR3 (**Fig. 1G**–**1H**). Enhancing mGluR3 levels did not improve memory performance in males. Of note, in mice without Cre expression, hippocampus-targeted injections of AAV vectors did not affect novelty exploration, anxiety-linked behavior, or learning and memory (**Fig. S8**), indicating that intrahippocampal delivery of AAV vectors was not sufficient to alter behavior in males or females.

These data suggest that aging- or disease-associated reductions of astrocytic mGluR3 expression may cause sex-specific neurocognitive outcomes and promote a divergence in disease onset or progression across sexes. However, mGluR3 is one of many types of G_i/o_-coupled receptors expressed by astrocytes. Consistent with our findings that astrocytic mGluR3 modulates memory, previous studies have shown that modulation of other non-glutamatergic astrocytic G_i/o_-coupled receptors (*5, 6*) or chemogenetic stimulation of astrocytic G_i/o_-coupled signaling can affect memory in males (*8, 9*). However, sex-dependence of these effects is not known because previous studies did not include females (*8, 9*). Thus, we used chemogenetics to acutely and non-invasively activate astrocytic G_i/o_-coupled signaling and test whether the sex-dependent effects of mGluR3 are unique to this receptor or reflect a broader feature of astrocytic G_i/o_-coupled receptors.

We generated an AAV vector encoding Cre-dependent HA-tagged hM4Di, a G_i/o_-coupled designer receptor exclusively activated by designer drugs (DREADD) (*51*), under the control of the *hGfaABC*_*1*_*D* promoter. mGluR3 and other G_i/o_-coupled receptors affect multiple intracellular signaling cascades, including Akt activation (**Fig. S9**), but cause minimal changes in calcium flux in hippocampal astrocytes (*25, 52-54*). Indeed, stimulation of hM4Di-expressing astrocytes with clozapine-N-oxide (CNO), the synthetic agonist for muscarinic DREADDs, increased the levels of phosphorylated Akt (**Fig. S10A**), indicating that the vector produced functional receptors at the cell surface. The hM4Di vector was delivered to the dorsal hippocampus of 4–7**-** month**-**old *Aldh1l1*-Cre mice, which induced astrocyte-selective and robust hM4Di expression in the hippocampus of male and female mice (**Figs. 3A, S10B**–**S10C**). Coverage and intensity of hM4Di protein expression in hippocampal subregions were similar between sexes (**Figs. 3A, S10B**). Pharmacokinetic data indicated that brain drug levels (clozapine) peaked 1 h after *in vivo* administration (**Fig. S10D**), which informed our study design. Thus, to test if biological sex influences the effects of astrocytic G_i/o_**-**coupled receptor activation on memory, *Aldh1l1***-**Cre mice expressing hM4Di in hippocampal astrocytes underwent training in the MWM 1 h after daily vehicle or CNO administration (**Fig. S10E**).

**Fig. 3.**
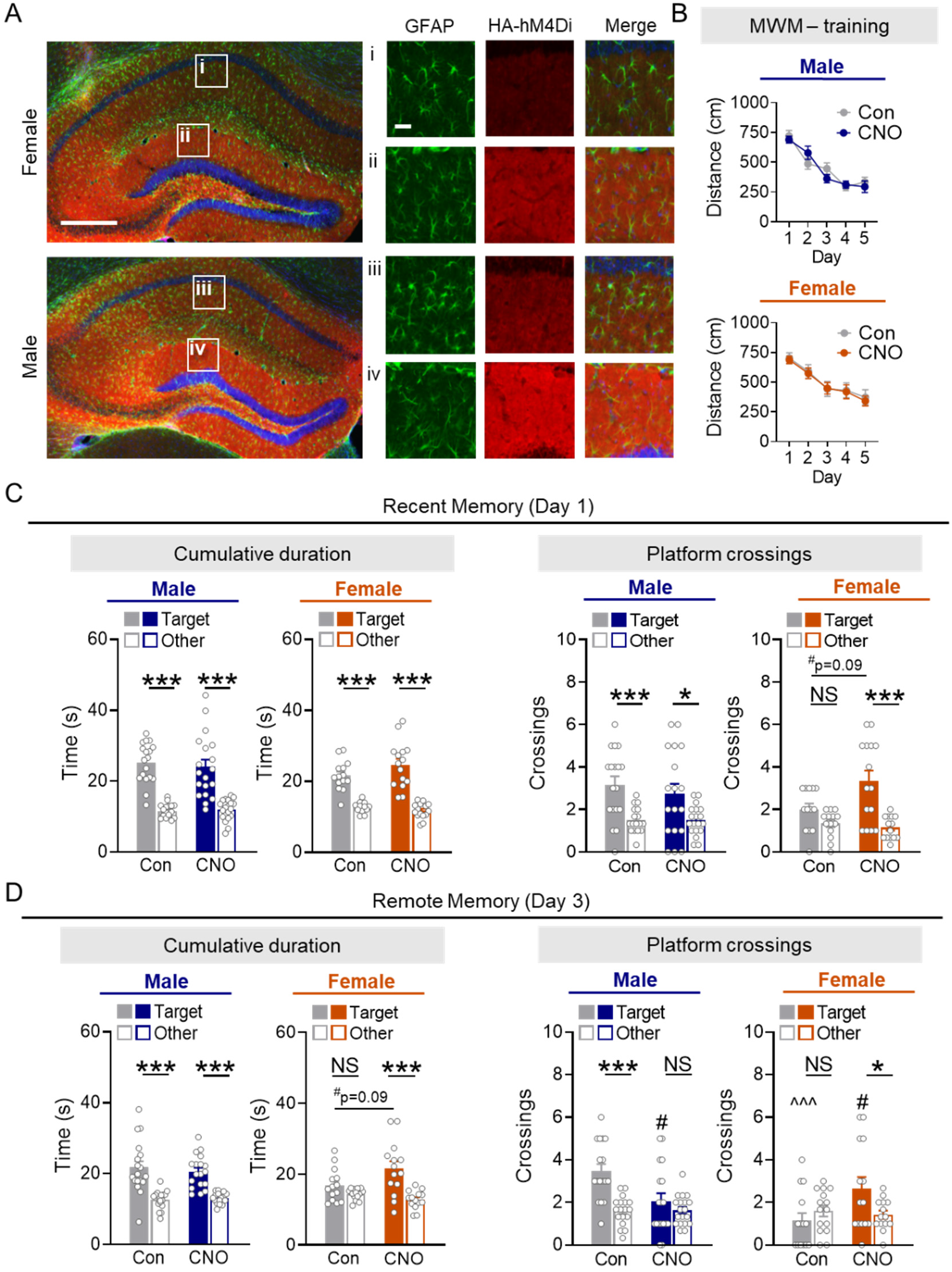
Chemogenetic stimulation of astrocytic G_i/o_-coupled signaling improves memory in females but impairs memory in males. (**A**) Representative images of GFAP (green) and HA-hM4Di (red) co-immunolabeling in hippocampal sections from *Aldh1l1*-Cre males and females in the presence of AAVs encoding HA-tagged hM4Di. Scale bars: 400 μm, 40 μm (insets). (**B**) hM4Di-expressing mice were trained in the Morris water maze. Data are shown as mean distance traveled per day. (**C** and **D**) Probe trials were performed one (C) or three (D) days after training. Two-way ANOVA (target duration, C): *F*(1, 62) = 1.540, p = 0.2193 for interaction effect. Two-way ANOVA (target crossings, C): *F*(1, 60) = 4.103, p = 0.0473 for interaction effect. Two-way ANOVA (target duration, D): *F*(1, 60) = 3.557, p = 0.0641 for interaction effect. Two-way ANOVA (target crossings, D): *F*(1, 59) = 12.81, p < 0.001 for interaction effect. Three-way ANOVA (B) or two-way ANOVA (C–D) with Sidak’s multiple comparisons post-hoc tests (vs control/sex): #p < 0.05; post-hoc tests (vs male target/condition): ^p < 0.001. Target preference was determined using Student’s t-test (C–D). *p < 0.05, ***p < 0.001. NS: no significant preference for target

Consistent with the effects of increasing astrocytic mGluR3 expression, hM4Di stimulation during training did not affect learning, but induced sex-specific effects on memory (**Fig. 3B**–**3D**). Specifically, astrocytic hM4Di stimulation improved recent (day 1) and remote (day 3) memory in females (**Fig. 3C**–**3D**), akin to the effects of increasing mGluR3 (**Fig. 2F**–**2G**), but impaired remote memory in males (**Fig. 3D**), which is consistent with recent findings in male mice (*8, 9*). These results suggest that astrocytic G_i/o_-coupled receptors as a class have the potential to exert sex-specific control of hippocampus-dependent behavior. Given that G_i/o_-coupled receptors are expressed by astrocytes throughout the brain, other brain regions may be subject to similar sex-specific modulation by astrocytes.

Notably, G_i/o_-coupled receptor signaling has convergent and antagonistic interactions with G_s_-coupled signaling (*53, 55*) (**Fig. S9**). Specifically, G_i/o_-coupled receptors, like mGluR3, inhibit cAMP synthesis, whereas G_s_-coupled receptors induce astrocytic cAMP synthesis and related signal transduction (*4, 53, 54*) and can also modulate G_i/o_-coupled signaling cascades (*56, 57*). Thus, to further examine which receptor mechanisms can promote sex-specific effects on memory, we next stimulated astrocytic G_s_-coupled receptors and assessed if their effects are in opposition to the effects observed with G_i/o_-coupled receptor stimulation. Similar to the hM4Di approach, we used an AAV vector encoding a Cre-dependent, HA-tagged G_s_-coupled receptor (rM3Ds) (*58*) under the control of *hGfaABC*_*1*_*D* promoter for astrocyte-selective expression. CNO treatment of rM3Ds-expressing astrocytes, but not control cells, increased the levels of phosphorylated CREB (**Fig. S11A**–**S11B**), confirming that astrocytic rM3Ds was intact and functional. Astrocytic expression of rM3Ds in the hippocampus of male and female *Aldh1l1*-Cre mice was consistent across hippocampal subregions and sexes (**Figs. 4A, S11C**) and was restricted to the hippocampus (**Fig. S11E**).

**Fig. 4.**
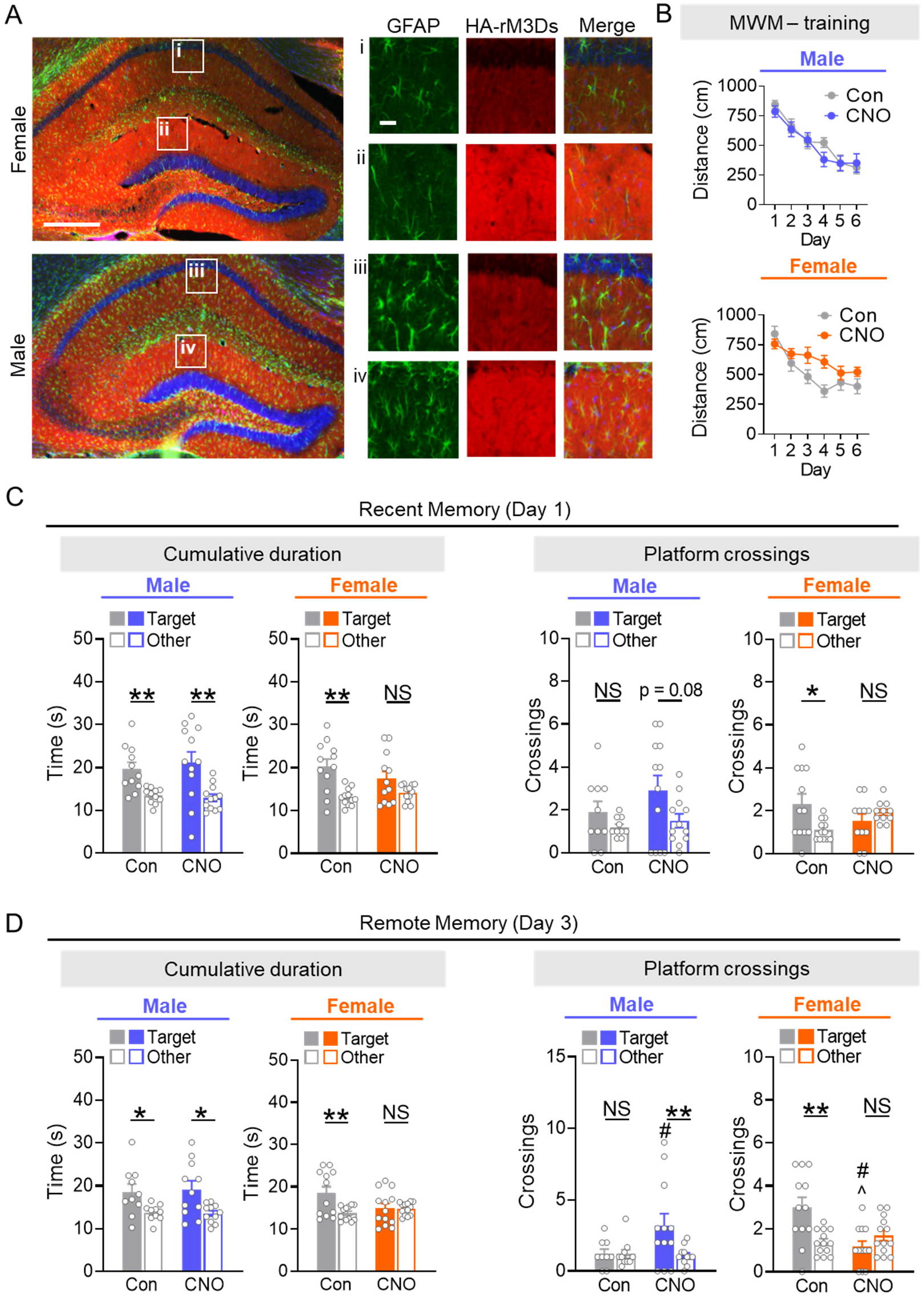
Chemogenetic stimulation of astrocytic G_s_-coupled signaling impairs memory in females but improves memory in males. (**A**) Representative images of co-immunolabeling for GFAP (green) and HA-rM3Ds (red) in mouse hippocampal sections from *Aldh1l1*-Cre males and females in the presence of AAVs encoding HA-tagged rM3Ds. Scale bar: 400 μm, 40 μm (insets). (**B**) rM3Ds-expressing mice were trained in the Morris water maze. Data are shown as mean distance traveled per day. (**C** and **D**) Probe trials performed one (C) or three (D) days after training. Three-way ANOVA (B) or two-way ANOVA (C–D) with Sidak’s multiple comparisons post-hoc tests (vs control/sex): #p < 0.05; post-hoc tests (vs male target/condition): ^p < 0.05. Target preference was determined using Student’s t-test (F–G). *p < 0.05, **p < 0.01. NS: no significant preference for target

Stimulation of astrocytic rM3Ds with daily CNO administration prior to MWM training (**Fig. S11D**) did not affect learning (**Fig. 4B)**, but impaired memory in females (**Fig. 4C**–**4D**) and enhanced remote memory in males (**Fig. 4D**). These consistent sex-specific effects on memory mirror the effects of mGluR3 knockdown (**Figs. 1G**–**1H**) and are opposite to the effects of increasing mGluR3 (**Fig. 2F**–**2G**) or stimulating hM4Di (**Figs. 3C**–**3D**). Our findings in males are also consistent with recent reports that optogenetic induction of cAMP synthesis in hippocampal astrocytes of males during training improves memory in a novel object recognition task (*59*), and that G_s_-coupled adrenergic receptors in hippocampal astrocytes facilitate memory consolidation in males (*3*). However, these studies did not test females and did not examine whether biological sex was a key determinant of the astrocytic effects on cognitive function.

Across different manipulations and GPCR subtypes, we found consistent patterns of divergent sex-specific effects of astrocytic receptors on memory (**Fig. S12**). These sex-dependent effects cannot be explained by potential behavioral or methodological artifacts in our different manipulations. Unlike mice used to alter mGluR3 levels, mice that underwent chemogenetic stimulation did not receive tamoxifen, an estrogen receptor modulator, to induce Cre recombinase activity, thus ruling out tamoxifen as a potential confound. Also, the effects on behavior cannot be attributed to off-target effects of CNO, because control mice that received AAV vectors encoding DREADDs but did not express Cre recombinase to enable DREADD expression were not responsive to CNO treatment (**Supplemental text** and **Fig. S13**).

Chemogenetic stimulation of G_s_-coupled or G_i/o_-coupled signaling did not affect swimming ability in either sex (**Fig. S14**). Furthermore, the effects of CNO on memory were diametrically opposite across the two different DREADDs and sexes (**Figs. 3**–**4**). Of note, the effects on memory in DREADD-expressing mice persisted for several days after CNO delivery was stopped. Our pharmacokinetic results indicate that CNO/clozapine is rapidly cleared from the brain within several hours (**Fig. S10D**), suggesting that astrocytic receptor activation during learning induced long-lasting changes related to memory consolidation and storage in both sexes.

Altogether, our findings suggest that astrocytic control of memory and related neural circuits involves a delicate balance between G_s_-coupled and G_i/o_-coupled receptor signaling and that dysregulation of this balance has opposing effects in males and females (**Fig. S12**). Our results reveal mGluR3 is one such endogenous astrocytic receptor with sex-specific roles in memory. Astrocytic GPCRs are linked to various molecular cascades through which astrocytes modulate glutamatergic (*29, 36, 53*) and GABAergic transmission (*55, 60*), highlighting that a broad spectrum of excitatory and inhibitory neural activities might be subject to sex-dependent regulation by astrocytic GPCRs. These findings raise important new questions regarding the roles of biological sex in astrocytic-neuronal interactions, including which sex-dependent molecular mechanisms are engaged by astrocytic GPCRs in males and females to influence memory, whether these mechanisms are established in early development or later in adulthood, whether they extend to astrocytes in other brain regions and to other neural functions and behaviors, and how these sex-dependent astrocytic-neuronal dynamics influence neurocognitive disorders in males and females.

Mechanistically, the observed sex-specific effects on memory may be mediated by intrinsic sex differences in astrocytic molecular pathways. Indeed, several astrocytic factors that can be regulated by GPCRs have been implicated in sex-specific effects, including cAMP response element-binding protein (CREB) (*61-63*), nuclear factor-kB (*64, 65*), and superoxide dismutase 2 (*66*). Astrocytic receptor signaling might also engage sex-specific neuronal mechanisms that result in distinct functional outcomes in a non-cell autonomous manner (*13, 16, 67, 68*). Further investigations are required to unravel if and how potential astrocyte-intrinsic and extrinsic neural circuit mechanisms mediate the divergent sex-specific effects of different astrocytic receptors, including mGluR3. Importantly, astrocytic GPCRs are altered in various neuropathologies (*4, 18, 26-28, 69-71*). Our findings suggest that these astrocytic perturbations may contribute to sex differences in cognitive impairments and that therapeutics that engage or perturb these receptors should be carefully assessed for sex-specific effects, a key consideration in personalized medicine (*17, 72*).

## Acknowledgments

We thank S. Tymchuk for technical support; M. Garvey, G. Coronas-Samano, L.J. Metakis, and E. Spencer for administrative support. We thank Drs. A. Rajadhyaksha, K. Pleil, O. Boudker, and L. Cantley for advice during project conceptualization and execution. We also thank Drs. T.A. Milner, A. Rajadhyaksha, K. Pleil, O. Boudker, A. Lee, Y. Xie, A. Krishnamurthy, and J. Fels for feedback on the manuscript. We thank B. Khakh for kindly providing *Aldh1l1*-CreERT BAC mice. All schematics were generated using BioRender.com.

## Author contributions

Conceptualization: SMM, ALM, ALO, AGO

Methodology: SMM, ALM, ALO, AGO

Investigation: SMM, FP, ALM, TSZ, DB, CZ, SRM, ALO, AGO

Formal Analysis: SMM, AGO Visualization: SMM, AGO Funding acquisition: SMM, AGO

Project administration: SMM, ALO, AGO Supervision: ALO, AGO

Writing – original draft: SMM, AGO

Writing – review and editing: SMM, ALM, FP, TSZ, DB, CZ, SRM, ALO, AGO

## Data and materials availability

Raw data are available from the corresponding authors upon request. All materials are available upon request or commercially.

## Supplementary Materials

Materials and Methods

Supplemental text

Figs. S1 to S14

Tables S1 to S3

