## Supplemental File for "Astrocytes regulate spatial memory in a sex-specific manner"

**This PDF file includes:**

Materials and Methods
Supplementary Text
Figs. S1 to S14
Tables S1 to S3

### Materials and Methods

Animals

All animal experiments were approved by the Institutional Animal Care and Use Committee of Weill Cornell Medicine and conducted in accordance with the guidelines set by the National Institutes of Health Guide for the Care and Use of Laboratory Animals. Mice were group-housed, with two to four mice per cage. Mice were maintained on a 12-h light/dark cycle with *ad libitum* access to food and water. All experiments were conducted during the light cycle and included littermate controls. In consideration of sex as a biological variable, all experiments were conducted to power each treatment group for both sexes at 80% power, with α set at 0.05. Although circulating hormone levels and estrous cycles were not monitored, we did not observe segmentation in the data and effects were typically consistent across multiple probe trials.

Astrocyte-targeted knockdown of mGluR3 was achieved using *Aldh1l1*-CreERT2 BAC transgenic mice crossed with Rosa26-LSL-Cas9-eGFP knock-in mice (*Aldh1l1*-CreERT2 x Rosa26-LSL-Cas9-eGFP). *Aldh1l1*-CreERT2 mice (Jackson Laboratory; strain #031008) express tamoxifen (TAM)-inducible Cre recombinase under the control of the astrocytic aldehyde dehydrogenase 1 family member L1 (*Aldh1l1*) promoter (kindly provided by Dr. Baljit S. Khakh). Rosa26-LSL-Cas9 mice (Jackson Laboratory; strain #026556) have Cre recombinase-dependent expression of the CRISPR-associated protein 9 (Cas9) endonuclease and eGFP under the CAG promoter. Use of Rosa26-LSL-Cas9-eGFP mice in combination with sgRNA vectors and a source for Cre recombinase enables manipulation of gene expression. The resulting *Aldh1l1*-CreERT2 x Rosa26-LSL-Cas9-eGFP cross enabled induction of Cas9 and eGFP expression selectively in astrocytes. Mice did not express Cas9-eGFP until treatment with TAM.

Astrocyte-targeted enhancement of mGluR3 expression was achieved using singly transgenic *Aldh1l1*-CreERT2 mice. One to two weeks post-stereotaxic injections, mice received single daily injections of TAM (Sigma, 80 mg/kg, s.c.) for five consecutive days. Behavioral testing began no sooner than five weeks after the completion of TAM treatment.

The expression of chemogenetic receptors hM4Di and rM3Ds was targeted to astrocytes using *Aldh1l1*-Cre mice (B6;FVB-Tg(Aldh1l1-cre)JD1884Htz/J). *Aldh1l1*-Cre mice were obtained from the Jackson Laboratory (strain #023748) and backcrossed for five generations onto the C57Bl/6J background. Similar to *Aldh1l1*-CreERT2 mice, *Aldh1l1*-Cre mice express Cre recombinase downstream of the astrocytic *Aldh1l1* promoter, but there is no dependence on TAM. Behavioral studies were conducted 3–5 weeks after stereotaxic injections.

Adeno-associated viral vectors (AAVs)

To knock down mouse mGluR3, AAV vectors were generated by removing the human synapsin promoter and mCherry sequences from pAAV-U6-sgRNA-hSyn-mCherry vector (donated by Dr. Alex Hewitt, Addgene #87916; RRID #Addgene 87916) (*73*) using the Anza 32 ApaI and Anza 14 SalI (Thermo Fisher Scientific) and the plasmid backbone was ligated with Anza T4 DNA Ligase Mix to create pAAV-U6-sgRNA. The existing 20 bp sgRNA spacer sequence was removed using Anza 33 LguI. sgRNAs targeting mouse *Grm3* were designed (*74*) and ordered as pairs of complementary unphosphorylated oligos (Integrated DNA Technologies):

#552 forward, CACCGCAGAGGTATCCAACGCCTGG;
#552 reverse, AAACCCAGGCGTTGGATACCTCTGC;
#703 forward, CCGCAGAGCATCGTTGACTAAAG;
#703 reverse, AACCTTTAGTCAACGATGCTCTGC.

Oligo pairs were phosphorylated, annealed, and ligated into the LguI-digested backbone to generate two new AAV vectors.

To enhance mGluR3 expression levels, the truncated human astrocyte-specific *GfaABC1D* promoter was digested from pAAV-GFAP-EGFP (donated by Dr. Bryan Roth, Addgene #50473; RRID #Addgene_50473) and cloned into pAAV-EF1a-DIO-hM4D(Gi)-mCherry (donated by Dr. Bryan Roth, Addgene #50461; RRID #Addgene_5046) using Anza 28 Mlul and Anza 14 SalI to replace the EF1a promoter. Rat mGluR3 with an N-terminal HA tag was synthesized as a gene block (Integrated DNA Technologies) and ligated in place of the hM4D(Gi) sequence of pAAV-*GfaABC1D*-DIO-hM4D(Gi)-mCherry using Anza 6 NheI and Anza 21 SgsI. To reduce construct size, the human growth hormone (hGH) poly-adenylation (polyA) sequence was replaced with the shorter bovine growth hormone (bGH) polyA sequence PCR amplified (Thermofisher Platinum SuperFi DNA Polymerase kit, Kit#12351002) from pCRII-TOPO CMV-cGFP-bGH Poly(A) (donated by Dr. Phil Sharp, Addgene #46835; RRID #Addgene_46835) (*75*) using primers containing XhoI and CpoI restriction sites. The resulting construct enabled Cre-dependent expression of mGluR3 in astrocytes.

For chemogenetic experiments, the $G_{i/o}$-coupled chemogenetic receptor hM4Di was PCR amplified from pAAV-GFAP-HA-hM4D(Gi)-IRES-mCitrine (donated by Dr. Bryan Roth, Addgene #50471; RRID #Addgene_50471) using primers containing NheI and SgsI restriction sites in order to replace the HA-mGluR3 sequence in the pAAV-*GfaABC1D*-DIO-HA-mGluR3 construct to generate pAAV-*GfaABC1D*-DIO-HA-hM4D(Gi) for Cre-dependent expression of HA-tagged hM4Di in astrocytes. Similar methods were employed to amplify the $G_s$-coupled chemogenetic receptor rM3D(Gs) from pAAV-GFAP-HA-rM3D(Gs)-IRES-mCitrine (donated by Dr. Bryan Roth, Addgene Plasmid #50472; RRID #Addgene_50472) to generate pAAV-*GfaABC1D*-DIO-HA-rM3D(Gs) for Cre-dependent expression of HA-tagged rM3Ds in astrocytes.

All vectors were transformed into NEB 5-alpha competent bacteria (New England Biolabs #C2987H) and purified with the Plasmid Plus Maxi Prep kit (Qiagen #12963). The integrity and accuracy of each AAV vector, including both AAV-ITRs, were verified by restriction digests and Sanger sequencing (Genewiz and Azenta) prior to viral production. AAV2/PHP.eB viral particles were produced by the Stanford University Gene Vector and Virus Core (sgRNA vectors) and the University of Pennsylvania (mGluR3, hM4Di, and rM3Ds). PHP.eB capsid vectors were provided courtesy of Drs. Viviana Gradinaru and Benjamin Deverman at the California Institute of Technology (*34, 35*). PHP capsids are a modification of AAV9 provided by the University of Pennsylvania.

Stereotaxic surgeries

Prior to surgery, mice were anesthetized with Avertin (2,2,2-tribromoethanol; Fisher Scientific #AC421430100, 200 mg/kg, i.p.) and supplemented with inhalant isoflurane (1%, Covetrus). During surgeries, meloxicam (2 mg/kg, s.c., Covetrus) and bupivacaine (1 mg/kg, t.d., Hospira) were administered for pain management. Once anesthetized, mice were secured in a stereotaxic frame (Kopf Instruments) and bilateral 1 mm-diameter openings were made in the skull using a mounted drill (Kopf Instruments). A 10 µl Hamilton syringe mounted to the stereotaxic frame was used to infuse 1 µl of sgRNAs (1:1 mixture of AAV2/PHP.eB-U6-sgRNA552 ($4.09 \times 10^{12}$ viral genomes (Vg)/ml) and AAV2/PHP.eB-U6-sgRNA703 ($2.11 \times 10^{12}$ Vg/ml)), or 0.5 µl AAV2/PHP.eB-hGfaABC1D-DIO-HA-mGluR3 ($8.15 \times 10^{12}$ Vg/ml), or 0.5 µl AAV2/PHP.eB-hGfaABC1D-DIO-HA-hM4Di ($1.12 \times 10^{13}$ Vg/ml), or 0.5 µl AAV2/PHP.eB-hGfaABC1D-DIO-HA-rM3Ds ($1.83 \times 10^{13}$ Vg/ml) into the dorsal hippocampus at the following stereotaxic coordinates (relative to bregma): -2.3 anterior/posterior; +/-2.0 medial/lateral; -2.0 dorsal/ventral. All AAV vectors were injected using a Micro 4 Microsyringe Pump (World Precision Instruments) at a rate of 0.1 µl/min. After each injection, the needle was left in place for an additional 5 min to allow for diffusion of the AAV. After needle withdrawal, the surgical site was sealed with Vetbond tissue adhesive (3M), and mice were continuously monitored under a heating lamp until fully recovered.

Behavioral testing

Age-matched littermates were distributed across experimental groups in home cages of each sex. Experimenters were blinded to experimental groups and mice were tested in random order. Before behavioral testing, except for the elevated plus maze, all mice were handled for 1–2 minutes per day for 5 days. Mice that were injured or in poor health were excluded from behavioral testing, regardless of experimental group. Prior to all behavioral tests, mice were acclimated to the testing room and lighting for 1 h. For experiments involving mGluR3 manipulations, behavioral testing began 5–6 weeks after completion of TAM administration. For chemogenetic experiments, behavioral testing began 3–5 weeks after stereotaxic injections.

*Elevated plus maze:* The elevated plus maze is a raised plus-shaped maze consisting of two enclosed arms and two open arms. The maze is raised 60–70 cm above the ground and placed under bright white light. After habituation to the room, the mice were placed in an open arm facing the center of the maze. Mice were allowed to freely explore the four arms of the plus-shaped maze for 5 minutes. The trial period was recorded and tracked using EthoVision XT video tracking software (Noldus Information Technology Inc., Leesburg VA). Between mice, the maze was cleaned with 70% ethanol to avoid interference between trials.

*Morris water maze:* Mice were tested in a 122 cm diameter pool filled with room temperature water (20 ± 2°C). The water was made opaque with white tempera paint (Colorations Non-Toxic Liquid Paint). Prior to testing, geometric spatial cues were set up around the pool. On the first day, mice underwent one session of three pre-training trials wherein mice were placed at one end of a rectangular channel (15 cm x 122 cm) with a square platform (14 x 14 cm) submerged 0.5 cm below the surface of the water at the other end of the channel. If a mouse did not swim to the channel within 15 s, the mouse was guided onto the platform, where it remained for 10 s before being gently dried and returned to the home cage. Three days following pre-training, the mice underwent hidden platform training in the full circular water maze.

During training, the platform was submerged approximately 1.0 cm below the surface of the water. Training consisted of one session per day for 5–7 consecutive days. In each training session, mice completed four training trials, wherein the drop location was varied but the platform location remained the same. The maximum trial time was 60 s. If the mouse did not find the platform within 60 s, it was gently guided to the platform by the experimenter. All mice sat on the target platform for 10 s after each training trial. During DREADD experiments, mice received injections of vehicle (0.5% DMSO in 0.9% saline, i.p.) or clozapine-N-oxide (CNO, Tocris #4936; 5 mg/kg, i.p.) 1 h prior to each training session.

During the probe trials, the hidden platform was removed from the pool and each mouse underwent one 60 s trial on each probe day, with the drop location kept constant at 180° angle from the target platform location. At the end of each trial, mice were removed from the pool, dried off to minimize stress, and placed back in the home cage.

Environmental or handling-related stress is known to affect behavioral readouts. Unlike mice that underwent mGluR3 manipulations, the mice used in chemogenetic experiments underwent daily intraperitoneal injections 1 h prior to training, which can cause stress and thus impair learning, accelerate the rate of forgetting, and affect remote probe performance in all groups (*76-78*). Thus, for chemogenetic experiments, we performed probe trials at three and six days post-training rather than nine days post-training. Similar sex-specific effects of hM4Di and rM3Ds stimulation were observed in probe trials at 6 days post-training (data not shown).

Rtrack analyses

Raw data were exported from Ethovision XT and processed and analyzed using R (version 4.0.3). Each individual trial was truncated to include only the first 20 s of the probe trial as previous studies have reported that this initial search period is the most sensitive to memory deficits (*40, 79*). The *Rtrack* package (version 1.0.0) (*41*) was used to determine the confidence score (percent match as a measurement of how well the swim path fit each model) for nine predefined swim strategies: non-goal-oriented strategies (thigmotaxis, circling, random), procedural strategies (scanning, chaining), and allocentric strategies (goal-directed search, corrected search, direct path, perseverance). Rtrack designers recommend a cutoff threshold of 0.4 for calling strategy (*41*). To maximize data inclusion, we did not set a threshold for confidence scores. Confidence scores ranged from 0.276–0.744 (64% over 0.4) for probe data in the sgRNA experiment, and from 0.218–0.80 (84% over 0.4) for probe data in the mGluR3 AAV experiment. Data are shown as the percentage of mice that utilized each swim strategy during a trial.

*Rtrack* was also used to calculate the initial trajectory error in each probe trial. This calculation assesses the distance of the mouse from the center of the goal (target platform) after a total swim distance equivalent to the calculated distance of a direct swim path from the starting point to the goal. This measure serves as a measure of the accuracy of the initial swim path.

Immunohistochemistry

Mice were anesthetized with Avertin (250 mg/kg, i.p.) and transcardially perfused for 3 min with 0.9% saline. Brains were removed and drop-fixed in PBS-buffered 4% paraformaldehyde (PFA) for 24 h, then rinsed three times with PBS before equilibrating with PBS-buffered 30% sucrose at 4°C until sectioning (at least 24 h). Mouse brains were sectioned at a thickness of 30 μm using a SM2010 R sliding microtome (Leica) equipped with a BFS-3MP freeing stage and cooling unit (Physitemp, Clifton, NJ). Free-floating sections were collected and stored in cryopreservative (30% ethylene glycol, 30% glycerol in PBS) for long-term storage at -20°C.

For all immunolabeling, free-floating sections were rinsed in PBS and permeabilized in PBS containing 0.5% Triton X-100 (PBS-T). Sections were blocked with 6% goat and/or donkey serum (Jackson ImmunoResearch) in PBS-T for 1–2 h at room temperature. Blocked sections were rinsed in PBS and incubated for 24 h (or for 48 h for Iba1 labeling) at 4°C in mouse anti-HA (1:500, 901513, Biolegend, RRID#AB_2565335), rabbit anti-GFAP (1:1,000, G9269, Sigma, RRID#AB_477035), mouse anti-GFAP (1:1,000, MAB3402B, Millipore, RRID#AB_10917109), rabbit anti-mGluR3 (1:500, ab166608, Abcam, RRID#AB_2833092), chicken anti-GFP (1:500, ab13970, Abcam, RRID#AB_300798), rabbit anti-NeuN (1:1000, ABN78, Millipore Sigma, RRID#AB_10807945), goat anti-NeuN (1:2500, ABN90, Millipore Sigma, RRID#AB_11205592), or goat anti-Iba1 (1:400, ab5076, Abcam, RRID#AB_222402 diluted in 3% serum in PBS-T.

Sections were rinsed and incubated for 1–2 h at room temperature in AlexaFluor-conjugated secondary antibodies (ThermoFisher Scientific #A-31573, A-31572, A-31571, A-31570, A-21432, A-21206, A-21202, RRID#AB_2536183, AB_162543, AB_162542, AB_2536180, AB_2535853, AB_2535792, AB_141607; Abcam ab150169, RRID# 2636803) diluted 1:500 in 3% serum in PBS-T before final rinses with PBS. Sections were mounted onto Superfrost slides (VWR), and coverslips were mounted with Vectashield (Vector Labs) or Prolong Diamond media (ThermoFisher Scientific).

Images were acquired using a BX710 microscope (Keyence) with a 20X objective (Nikon). Images were stitched with BZ-X Analyzer Software (Keyence) and analyzed with ImageJ (FIJI). For high-resolution images of mGluR3 and HA labeling, images were captured using an LSM880 confocal microscope (Zeiss) equipped with a 40X objective (Zeiss) and Zen Black v2.3 SP1 FP3 acquisition software (Zeiss). Images were processed in FIJI to create Z-stack overlays of representative fields.

Microfluidic RT-qPCR

Saline-perfused brain tissue was flash frozen in isopentane on dry ice and stored at -80°C until dissection. Briefly, frozen hemibrains were affixed in warm 30% low-melting point agarose and rapidly cut to 450 µm sections using the McIlwain Tissue Chopper (Stoelting #51350). The sections were quickly placed in ice-cold PBS and the dentate gyrus and the hippocampal CA1 region were dissected under a dissecting microscope (AmScope), placed in RNase-free low-binding tubes (Eppendorf), and re-frozen on dry ice until storage at -80°C. Dissected tissue was homogenized in extraction buffer (Qiagen RLT with β-mercaptoethanol) in a pre-chilled adaptor tube rack using a Fisherbrand Bead Mill 24 (Fisher Scientific #15-340-163) for 20 s with speed setting 5.

RNA was purified using the RNeasy Mini Kit with on-column DNase treatment following manufacturer's instructions (Qiagen #74106, #79256), denatured for 5 min at 70°C and reverse transcribed with the Protoscript First Strand Synthesis Kit (New England Biolabs #6300L). cDNA was pre-amplified for 14 cycles against a pool of selected primers (Supplemental Table 3) using PreAmp Grandmaster mix (TATAA Biocenter, Sweden #TA05). Pre-amplified DNA underwent exonuclease I treatment (New England Biolabs #M0293L) before being diluted 10-fold with nuclease-free water and mixed with SsoFast EvaGreen with Low ROX (BioRad #1725211). Samples were mixed with chip-specific reagents (Fluidigm) and then loaded into a 96.96 Dynamic Array Chip (Fluidigm #100-6308, BMK-M-96.96). The chip inlets were loaded with individual primers pre-mixed with DNA assay reagent (Fluidigm). The IFC Controller HX was used to prime and load the chip in preparation for qPCR. Amplification and melting curves were measured and analyzed with the BioMark HD System (Fluidigm). Cycle of quantification (Cq) values were thresholded by the BioMark software and normalized to the average of reference genes (*Actb*, *Gapdh*, *Gusb*, *Tbp*). Normalized Cq values were then used to determine the ddCq and fold change relative to control groups.

Primary astrocyte cultures

All cultures were maintained at 37°C in a humidified 5% $CO_2$-containing atmosphere. Cortices and hippocampi from postnatal day 3 mouse pups (non-transgenic or *Aldh1l1*-Cre) were cleared of meninges, dissociated by manual trituration with a P1000 pipette, and plated into culture plates pre-coated with poly-D-lysine (50 µg/ml, Sigma-Aldrich). Cells were maintained in DMEM supplemented with 20% heat-inactivated FBS (VWR #89510-188), 1X GlutaMAX (ThermoFisher #35050061) and 1 mM sodium pyruvate (Thermo Scientific). Cells were washed after 4–5 days in culture and maintained in fresh media after washing. After 9–11 days in culture, medium was replaced with DMEM without serum and glutamine, and cells were treated 18–24 h later with clozapine-N-oxide (CNO, Tocris #4936, 5 µM) or LY354740 (Tocris #3246, 1 µM) for 10 min prior to lysis to assess the levels of phosphorylated and total proteins by Western blotting.

Western blotting

Cultured astrocytes were rinsed with ice-cold PBS before aspirating buffer and lysing on ice for 10 min with ice-cold buffer containing 10 mM Tris (pH 7.4), 150 mM NaCl, 0.5% deoxycholate, 5 mM EDTA, 0.5% Triton X-100, protease inhibitor cocktail (Roche #11836153001), and two phosphatase inhibitor cocktails (Sigma-Aldrich P5726 and P0044). Harvested lysates were sonicated on ice for 5 s at 10% power with a probe sonifier (Branson), centrifuged at 10,000 rpm for 10 min at 4°C, and the supernatants assayed for protein content (Pierce Detergent-Compatible Bradford Assay, ThermoFisher #23246).

Samples (10–20 μg/well) were separated using 4–12% NuPAGE Bis-Tris gels (ThermoFisher #NP0336BOX) and transferred to nitrocellulose membranes using a Mini Blot Module (ThermoFisher). Membranes were blocked for 1 h in 5% bovine serum albumin (BSA) (VWR #97062-904) in Tris-buffered saline prior to incubation overnight at 4°C in 3% BSA in TBS containing 0.2% Tween-20 (TBS-T) and the following primary antibodies: rabbit anti-phospho-CREB (S133) (1:1000, Cell Signaling 9198, RRID#AB_2561044) and mouse anti-total-CREB (1:250, Cell Signaling 9104S, RRID#AB_490881), rabbit anti-phospho-Akt (Ser473) (1:2500, Abcam ab81283, RRID#AB_2224551) and mouse anti-total-Akt (1:250, Cell Signaling 2920S, RRID#AB_1147620), or rabbit anti-mGluR3 (1:500, Abcam ab166608, RRID#AB_2833092) and mouse anti-γ-tubulin (1:2500, Sigma T5326, RRID#AB_532292).

After washing, membranes were incubated in IR Dye 680RD donkey anti-mouse (1:15,000; LI-COR #926-68072, RRID#AB_2814912) and IR Dye 800CW donkey anti-rabbit (1:15,000; LI-COR #926-32213; RRID#AB_621848) in 3% BSA in TBS-T for one hour. Blots were rinsed twice with TBS-T and once with TBS and then dried before imaging on the Odyssey CLx scanner (Licor). Immunoblotting was quantified using LI-COR Image Studio software.

Pharmacokinetic analyses

Clozapine and CNO measurements in brain tissue were performed by Charles River Laboratories (South San Francisco, US). Analytes were extracted from pre-weighed hippocampal tissue using standard methods and quantified by LC-MS/MS against analyte calibration curves.

Statistical analyses

Statistical specifications are reported in the figures and corresponding figure legends. Numbers of mice used in each experiment are listed in Tables S1 and S2. All data are presented as mean ± S.E.M. Statistical tests were performed using GraphPad Prism 8, except Fisher's exact test, which was performed using IBM SPSS Statistics for Windows, Version 24.0. The criterion for data point exclusion was established during the design of the study and was set to values above or below two standard deviations from the group mean. For determination of target preference in the MWM, mice with thigmotaxis were also excluded. Two-sided Student's t tests were used to determine statistical significance between two groups. Welch's correction was used to account for unequal variances. Differences among multiple groups were assessed by one-way, two-way, or three-way ANOVAs followed by Sidak's multiple comparisons post-hoc tests, as specified in the legends. Null hypotheses were rejected at $p < 0.05$.

### Supplementary Text

<u>Imaging of endogenous and AAV vector-induced mGluR3 protein expression required different acquisition settings</u>

The levels of endogenous mGluR3 expression appear low in Fig. 2A–B, but this is a technical limitation due to the low exposure settings required to capture the higher levels of mGluR3 in AAV-injected mice without saturating the signal. We have provided additional images (Fig. S5) to demonstrate that endogenous mGluR3 expression is robust in control mice after adjusting the exposure settings.

<u>DREADD-independent effects of CNO do not account for the observed effects on memory</u>

Previous studies have shown that CNO is readily converted to clozapine, a psychoactive compound that stimulates endogenous GPCRs, including 5-$HT_{2A}$ and dopamine D2 receptors (*80*). Indeed, our own pharmacokinetic analyses demonstrate that clozapine, but not CNO, is detectable in the hippocampus for hours post-injection. Thus, in additional control experiments, we tested whether CNO administration induced DREADD-independent effects on memory in males or females. We found that AAV-injected non-transgenic mice did not have detectable DREADD expression (Fig. S13A), and CNO administration (5 mg/kg, i.p.) did not affect learning in males or females (Fig. S13B). Despite a trend for increased time in the target quadrant in CNO-treated females (Fig. S13C), CNO treatment did not affect probe crossings or performance at later timepoints (Fig. S13C–D) and could not account for the robust memory enhancement induced in females expressing hM4Di (Fig. 3C–D), the profound memory loss induced in females expressing rM3Ds (Fig. 4C–D), and the opposite CNO-induced effects on memory observed in male mice with DREADD expression (Figs. 3–4).

<u>Tamoxifen-induced effects do not account for the observed sex differences in memory</u>

Tamoxifen (TAM) is a known estrogen receptor modulator that can facilitate estrogen receptor degradation and act as a mixed agonist and antagonist. However, short-term TAM administration has not been shown to alter EPM or MWM performance in either sex (*81*). Nonetheless, we designed our experiments to minimize potential interactions between our manipulations and TAM administration. In *Aldh1l1*-CreERT2 mice with mGluR3 manipulations, all behavioral experiments were performed no less than 5 weeks after cessation of TAM administration. TAM and its psychoactive metabolites are minimally detectable by one week after administration (*82*). Thus, we conclude that there was minimal-to-no influence of TAM on the behavioral measurements in our study. Furthermore, the sex-specific effects observed with mGluR3 manipulations were recapitulated with chemogenetic stimulation using independent mouse lines and AAV vectors. The chemogenetic experiments were conducted in *Aldh1l1*-Cre mice that were not exposed to TAM.

<u>Additional computational analyses of spatial search strategies further reveal sex-specific effects of astrocytic mGluR3 on memory</u>

We performed computational analyses of MWM probe trial performance in mice with either reduced or increased astrocytic mGluR3 expression. We chose to perform these analyses on the first 20 s of each probe trial because this period typically corresponds to the most directed search for the platform location, whereas later periods of the probe trials are more akin to extinction (*40, 79*).

Astrocytic mGluR3 knockdown did not affect search strategies in males or females during recent probe trials (one day post-training), despite deficits in target preference in females (Fig. 1G–H). These results suggest that mGluR3 knockdown reduced the accuracy of spatial memory for the target location in females without significantly altering the overall search strategies (Fig. S3C) or trajectories (Fig. S3E). In contrast, males with mGluR3 knockdown had improved target preference (Fig. 1H) and significantly different search strategies in the remote probe trial relative to control males (9 days post-training; Fig. S3D). In males, mGluR3 knockdown increased the temporal durability of goal-directed search strategies (Fig. S3D) and reduced error in the search trajectories (Fig. S3F), further supporting the conclusion that mGluR3 knockdown improved spatial memory in male mice.

In mice with enhanced astrocytic mGluR3 expression, there were no differences detected in search strategies or trajectories in recent probe trials (one day post-training; Fig. S7C–S7F), in concurrence with the minimal changes observed in target preferences (Fig. 2F). However, in the remote probe trials (9 days post-training), females had significant differences in search strategies as compared to males. Aging control females used fewer goal-directed strategies than aging control males, consistent with the observed impairments in target crossings (Figs. S7C, S7D), further suggesting that older females develop spatial memory deficits at earlier ages than males. Notably, increasing astrocytic mGluR3 expression increased the percentage of female mice that used the goal-directed search strategy (Fig. S7D), without detectable changes in initial trajectory errors (Fig. S7F), further revealing improvements in memory performance in older females with enhanced astrocytic mGluR3 expression. No changes were detected in search strategies and trajectories used by male mice with increased mGluR3 expression (Fig. S7C, S7E).

These complementary analyses provide additional context for how astrocytic mGluR3 manipulations might modulate memory performance across sexes. In females, reductions in astrocytic mGluR3 impaired the accuracy of spatial memory without altering learning or overall search strategies. Together, these results implicate hippocampal place cells in the CA3 region, which have stable responses to familiar environments during memory retrieval (*83*). CA3 hyperactivity is associated with age-related cognitive deficits in rodents (*84, 85*) and humans (*86*). In rodents, attenuation of CA3 hyperactivity is sufficient to rescue age-related memory loss (*87*). Middle-aged female mice with lower mGluR3 expression compared to age-matched males had impaired spatial memory. Intriguingly, female mice at this age (10–12 mo) have increases in synaptophysin labeling in the CA3, whereas males do not have these changes until older ages (18–20 mo) (*88*). Thus, increasing astrocytic mGluR3 expression might have improved memory in females but not males at least in part by reversing age-related functional changes in the CA3.

In male mice with knockdown of astrocytic mGluR3, we observed improvements in target memory, search strategies, and trajectories in remote probe trials (Figs. 1H, S3D, S3F). Consistent with our results, activation of astrocytic $G_{i/o}$-coupled receptors in the CA1 region has recently been shown to impair remote memory in males, and this effect involved astrocytic modulation of CA3-CA1 neuronal activities and downstream projections to the anterior cingulate cortex (*8*). Thus, improvements in remote memory in males following knockdown of the $G_{i/o}$-coupled mGluR3 or stimulation of $G_s$-coupled receptors might involve enhanced functions in similar circuits.

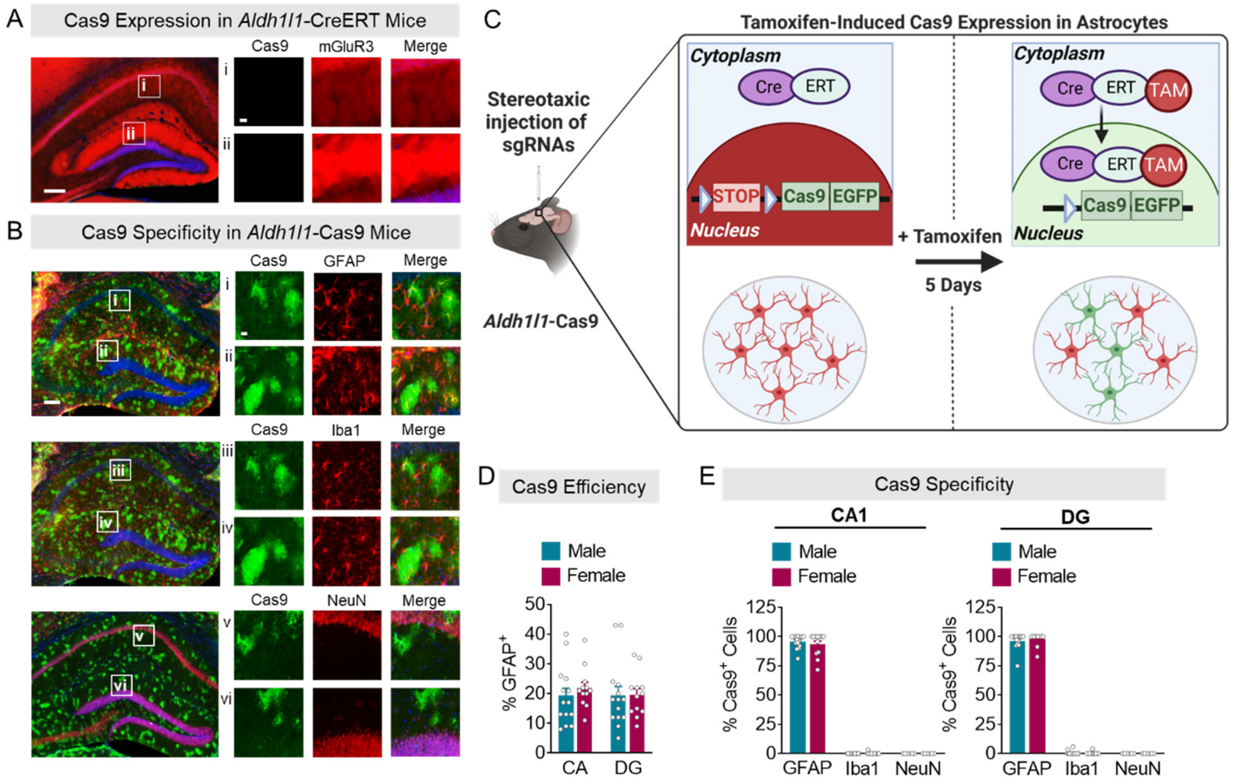


**Fig. S1. Further characterization of *Aldh1l1*-Cas9 mice.** (**A**) Representative images of mGluR3 (red) and Cas9-eGFP (green) co-immunolabeling from the CA1 stratum radiatum (CA1sr) (i) and DG molecular layer (DGmol) (ii) in sgRNA-injected *Aldh1l1*-CreERT2 single transgenic control mice. Scale bars: 400 µm, 40 µm (inset). (**B**) Representative images of Cas9-eGFP (green) co-immunolabeling with cell type-specific markers (red), including GFAP for astrocytes (i and ii), Iba1 for microglia/macrophages (iii and iv), and NeuN for neurons (v and vi) in *Aldh1l1*-Cas9 doubly transgenic mice. Scale bars: 400 µm, 40 µm (inset). (**C**) Schematic illustrating the inducible doubly transgenic *Aldh1l1*-Cas9 mouse model that is amenable to sgRNA-mediated gene knockdown in astrocytes. (**D**) Immunofluorescence-based quantification of the percentage of GFAP-positive cells that were also Cas9-eGFP-positive in the CA1sr and DGmol regions of the hippocampus of tamoxifen-injected *Aldh1l1*-Cas9 mice. (**E**) Immunofluorescence-based quantification of the percentage of Cas9-eGFP-positive cells that also expressed a cell type-specific marker for astrocytes (GFAP), microglia/macrophages (Iba1), or neurons (NeuN) in the CA and DG regions of the hippocampus of tamoxifen-injected *Aldh1l1*-Cas9 mice.

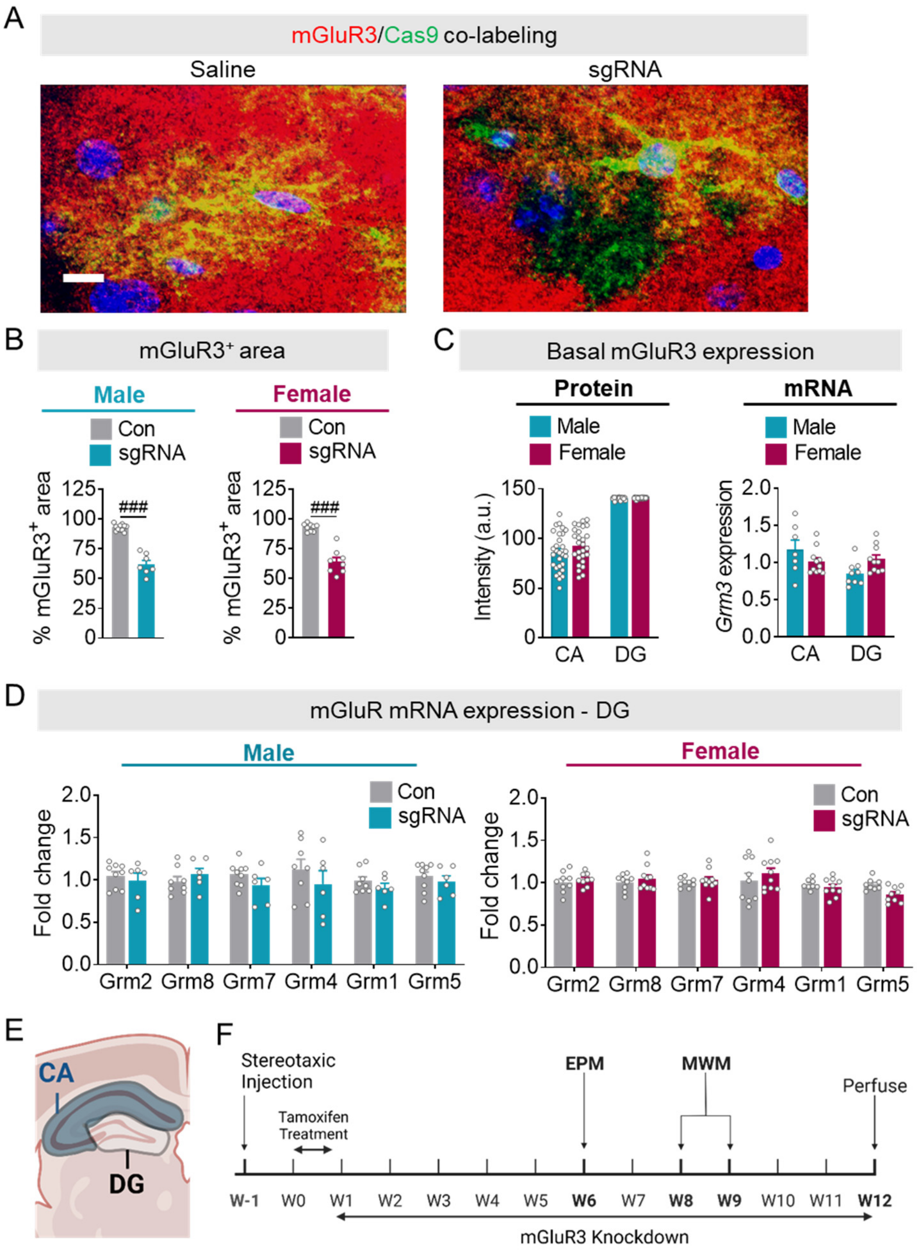


**Fig. S2. Further characterization of mGluR3 (*Grm3*) expression and the effects of sgRNAs.** (**A**) Representative images of mGluR3 (red) and Cas9-eGFP (green) co-immunolabeling in saline or sgRNA-injected *Aldh1l1*-Cas9 mice. Images highlight astrocytic loss of mGluR3 in Cas9-positive cells in the dentate gyrus (DG). Scale bar: 10 µm (**B**) Immunofluorescence-based quantification of the % DG area that was mGluR3-positive in saline-injected (Con) and sgRNA-injected *Aldh1l1*-Cas9 mice. Two-way ANOVA: $F(1, 31) = 39.40$, $p < 0.001$ for main effect of sgRNA; $F(1, 31) = 0.5896$, $p = 0.4484$ for interaction effect. (**C**) Immunofluorescence-based (Protein) and RT-qPCR-based (mRNA) quantification of mGluR3 levels in the CA and DG regions of saline-injected male and female mice. Two-way ANOVA (Protein): $F(1, 48) = 0.2759$, $p = 0.6018$ for main effect of sex, $F(1, 31) = 0.5896$, $p = 0.4484$ for interaction effect. Two-way ANOVA (mRNA): $F(1, 33) = 0.0327$, $p = 0.8576$ for main effect of sex, $F(1, 33) = 0.0162$, $p = 0.0162$ for interaction effect. (**D**) RNA levels of different mGluR subtypes in the DG of mice with or without mGluR3 knockdown, in order of sequence homology to mGluR3. Three-way ANOVA: $F(5, 173) = 1.012$, $p = 0.4122$ for interaction effect. (**E**) Schematic illustrating the hippocampal dissections for RT-qPCR. (**F**) Experimental timeline for mice with astrocytic mGluR3 knockdown. Two-way ANOVA (B–C) and three-way ANOVA (D) with Sidak's multiple comparisons post-hoc tests (vs control/sex): ###$p < 0.001$

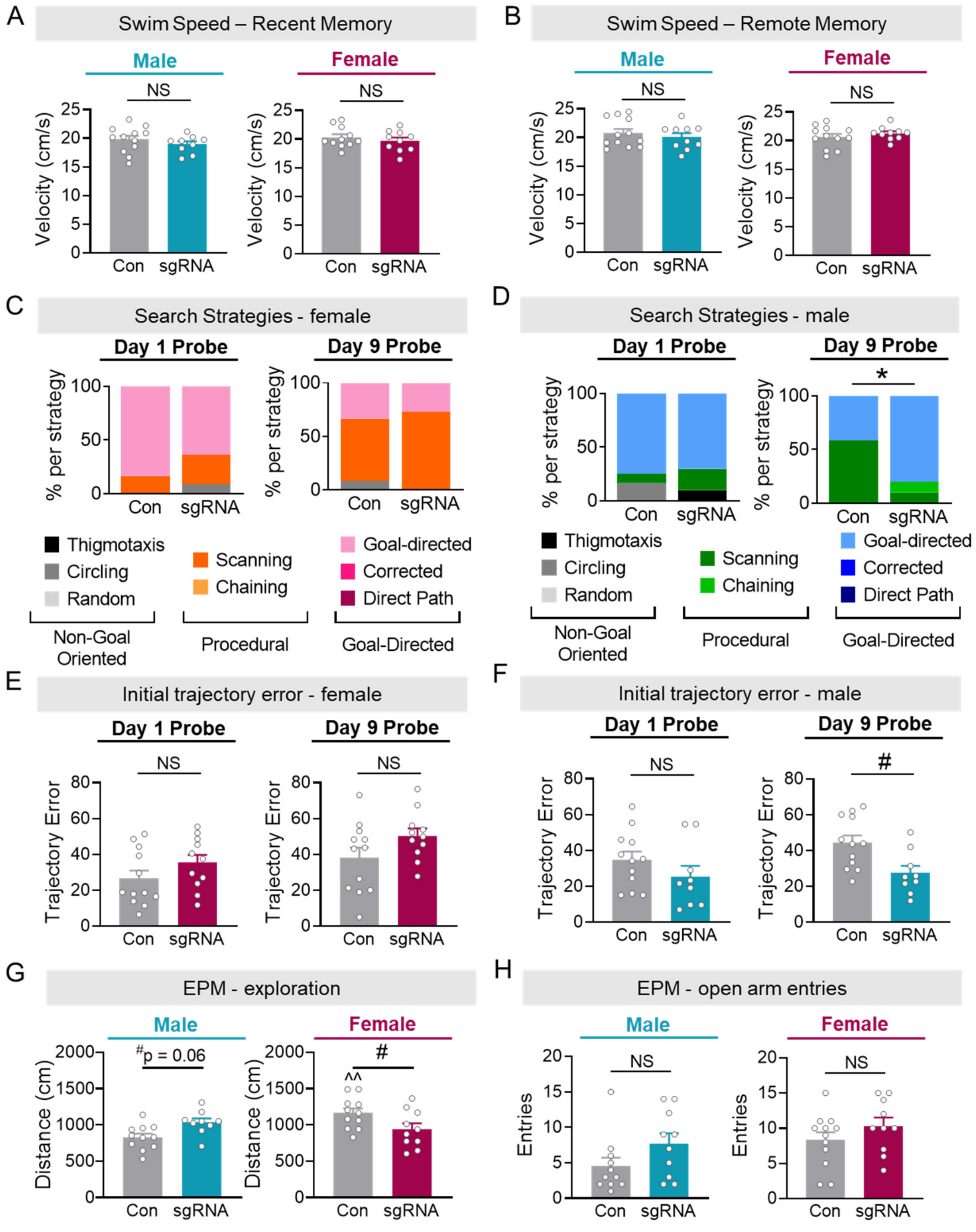


**Fig. S3. Additional behavioral characterization of *Aldh1l1*-Cas9 mice with or without mGluR3 knockdown in hippocampal astrocytes.** (**A** and **B**) Swim speeds in the Morris water maze during probe trials conducted one (A) and nine (B) days post-training. Two-way ANOVA: $F(1, 38) = 0.0350$, $p = 0.8525$ for interaction effect (A); $F(1, 40) = 1.167$, $p = 0.2866$ for interaction effect (B). (**C** and **D**) Search strategies in female (C) and male (D) mice during recent or remote probe trials quantified with Rtrack. (**E** and **F**) Initial trajectory errors in female (E) and male (F) mice during recent or remote probe trials quantified with Rtrack. (**G**) Exploration in the elevated plus maze (EPM). Two-way ANOVA: $F(1, 37) = 10.88$, $p = 0.0022$ for interaction effect. (**H**) Open-arm entries in the EPM. Two-way ANOVA: $F(1, 39) = 0.2292$, $p = 0.6348$ for interaction effect. Two-way ANOVA (A–B and F–H) with Sidak's multiple comparisons post-hoc tests (vs control/sex): #$p < 0.05$; post-hoc tests (vs male/condition): ^^$p < 0.01$, NS: not significant. Fisher's exact test (C–D): *$p < 0.05$.

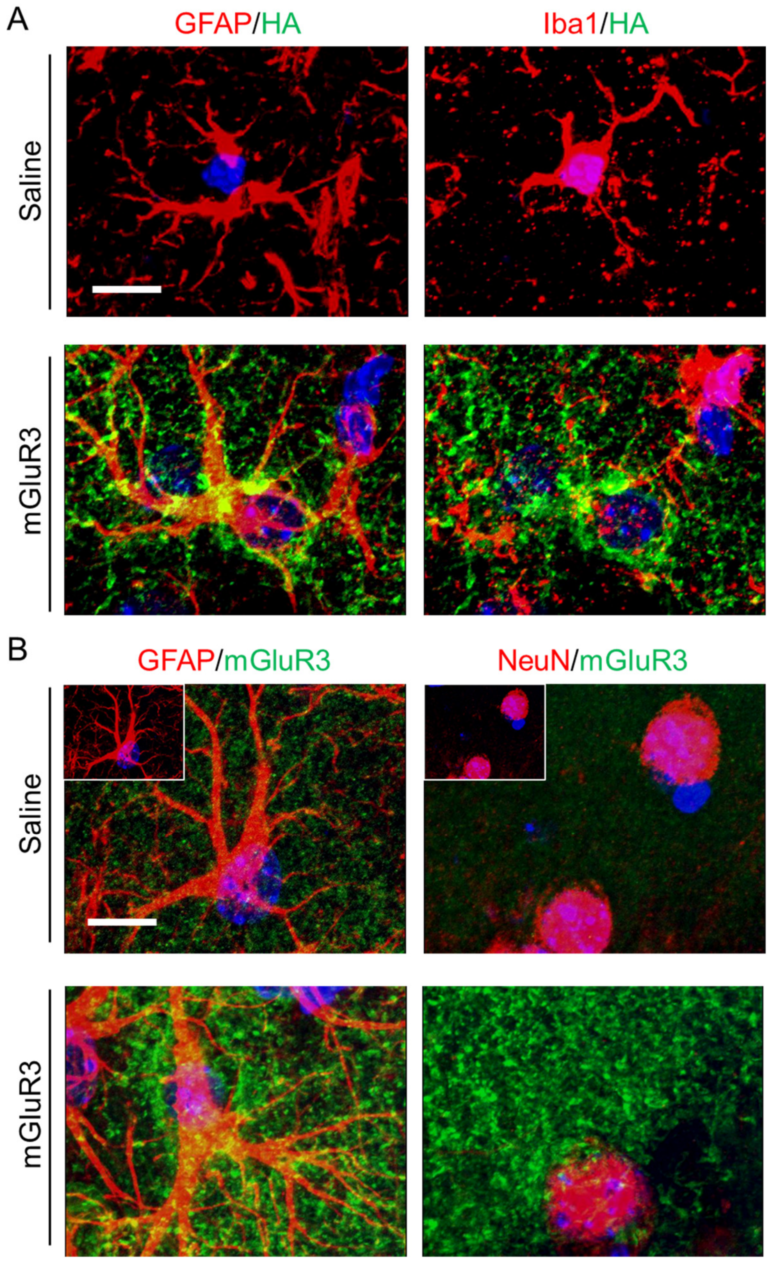


**Fig. S4. High-resolution images of hippocampal sections from *Aldh1l1*-CreERT2 mice injected with saline or AAV vector encoding HA-tagged mGluR3.** (**A**) Representative images of HA-mGluR3 expression (green) and GFAP or Iba1 (red) co-immunolabeling in the DG of *Aldh1l1*-CreERT2 mice injected with either saline or mGluR3-encoding AAV vector. Scale bar: 10 µm. (**B**) Representative images of total mGluR3 expression (green) and GFAP (astrocyte, red) or NeuN (neuron, red) co-immunolabeling in the DG of *Aldh1l1*-CreERT2 mice injected with either saline or mGluR3-encoding AAV vector. Differences in mGluR3 intensity required the use of different image acquisition and processing settings for the mGluR3 signal. Insets show images of saline-injected mice in which mGluR3 signal was processed similarly to mice with AAV injections. Scale bar: 10 µm.

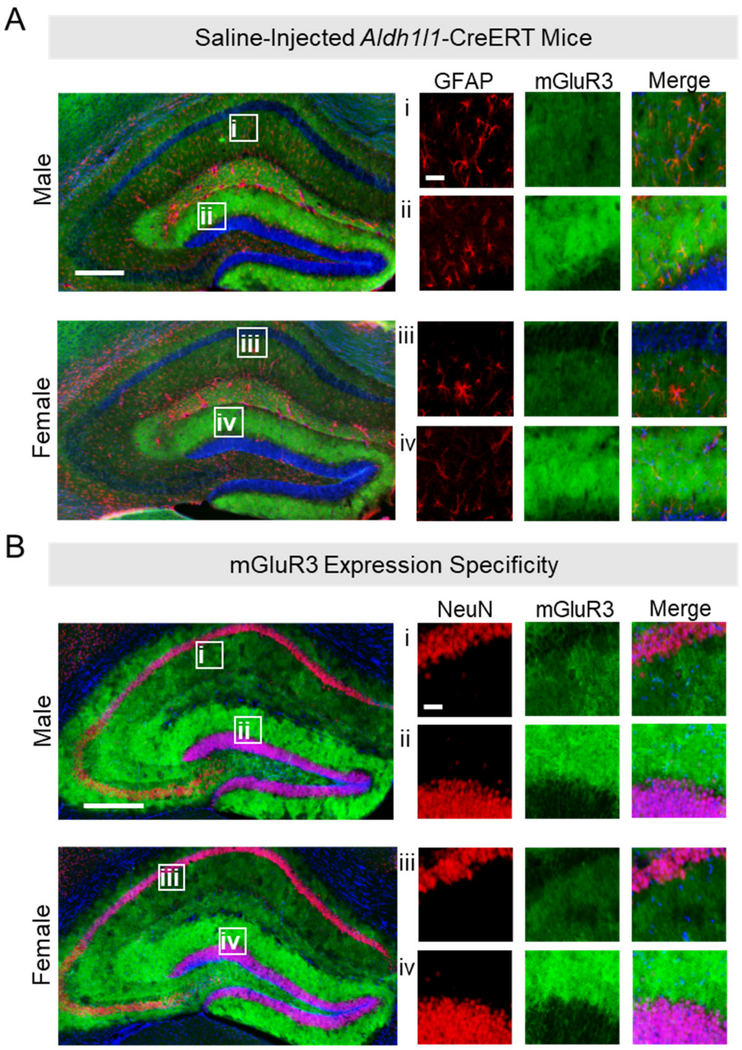


**Fig. S5. Additional characterization of the mGluR3-encoding AAV vector.** (**A**) Representative images of mGluR3 (green) and GFAP (red) co-immunolabeling in the CA1 and DG of saline-injected (control) *Aldh1l1*-CreERT2 mice. Scale bars: 400 µm, 40 µm (inset). (**B**) Representative images of mGluR3 (green) and NeuN (red) co-immunolabeling in the CA1 and DG of AAV vector-injected *Aldh1l1*-CreERT2 mice. Scale bars: 400 µm, 40 µm (inset).

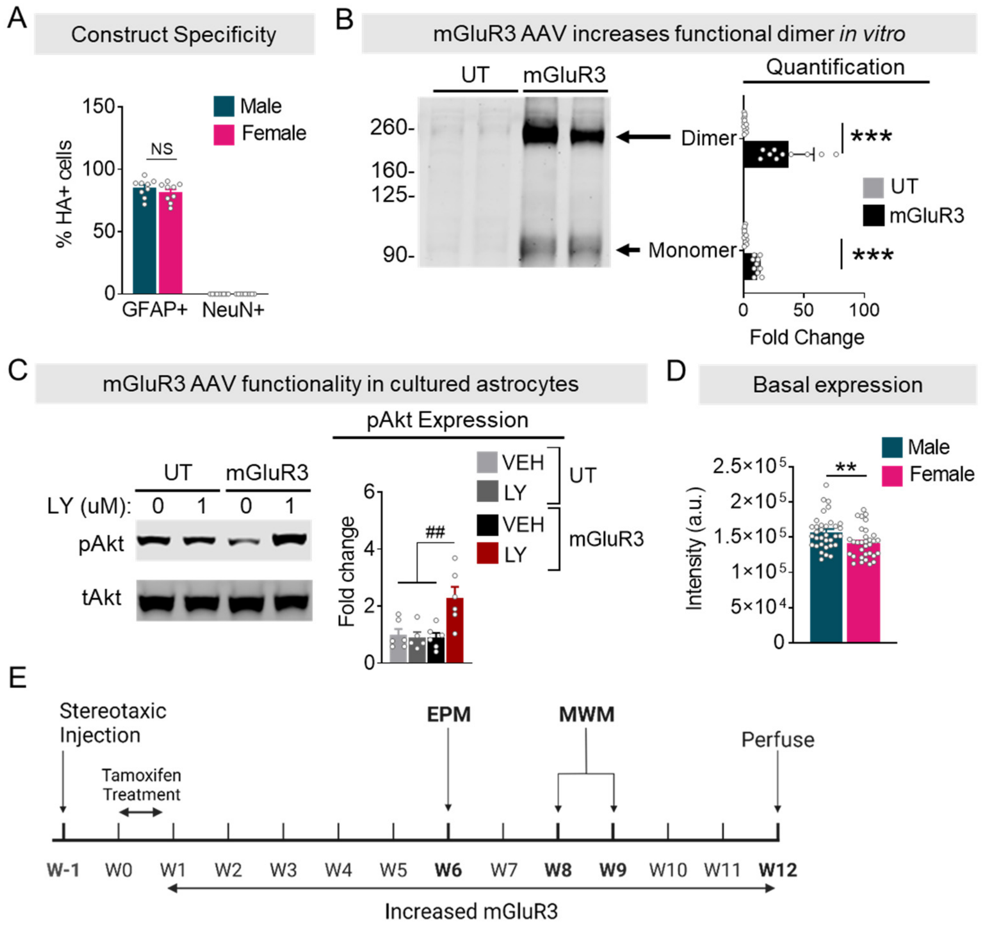


**Fig. S6. Validation of the mGluR3-encoding AAV vector.** (**A**) Immunofluorescence-based quantification of the percentage of GFAP-positive and NeuN-positive cells that were also HA-positive in the DG region of the hippocampus in male and female *Aldh1l1*-CreERT2 mice injected with the mGluR3-encoding AAV vector. Two-way ANOVA: $F(1, 32) = 2242$, $p < 0.001$ for main effect of cell marker; $F(1, 32) = 1.057$, $p = 0.311$ for main effect of the sex; $F(1, 39) = 16.66$, $p = 0.0002$ for the interaction. (**B**) Representative Western blot image and quantification of mGluR3 dimer and monomer levels in primary cultured astrocytes untransduced (UT) or transduced with a Cre-dependent AAV vector encoding mGluR3. Experiments were performed in two independent cultures. Two-way ANOVA: $F(1, 39) = 16.66$, $p = 0.0002$ for main effect of isoform; $F(1, 39) = 53.05$, $p < 0.0001$ for main effect of the AAV; $F(1, 39) = 16.66$, $p = 0.0002$ for the interaction. (**C**) Representative Western blot images and quantification of phospho-Akt and total Akt levels following mGluR3 stimulation with LY354740 (LY, 1 µM, 10 min) in cultured astrocytes that were untransduced (UT) or transduced with a Cre-dependent AAV vector encoding mGluR3. Experiments were performed in two independent cultures. Two-way ANOVA of pAkt: $F(1, 19) = 8.163$, $p = 0.0101$ for interaction effect. (**D**) Baseline mGluR3 immunofluorescence quantified in the DG of nontransgenic control mice injected with saline. (**E**) Experimental timeline for mice with increased mGluR3 expression. Data in (A–B) and (D) were analyzed with Student's t-tests. $^{**}p < 0.01$, $^{***}p < 0.001$. Data in (C) were analyzed with the two-way ANOVA and Sidak's post-hoc test. ##$p < 0.01$. NS = not significant.

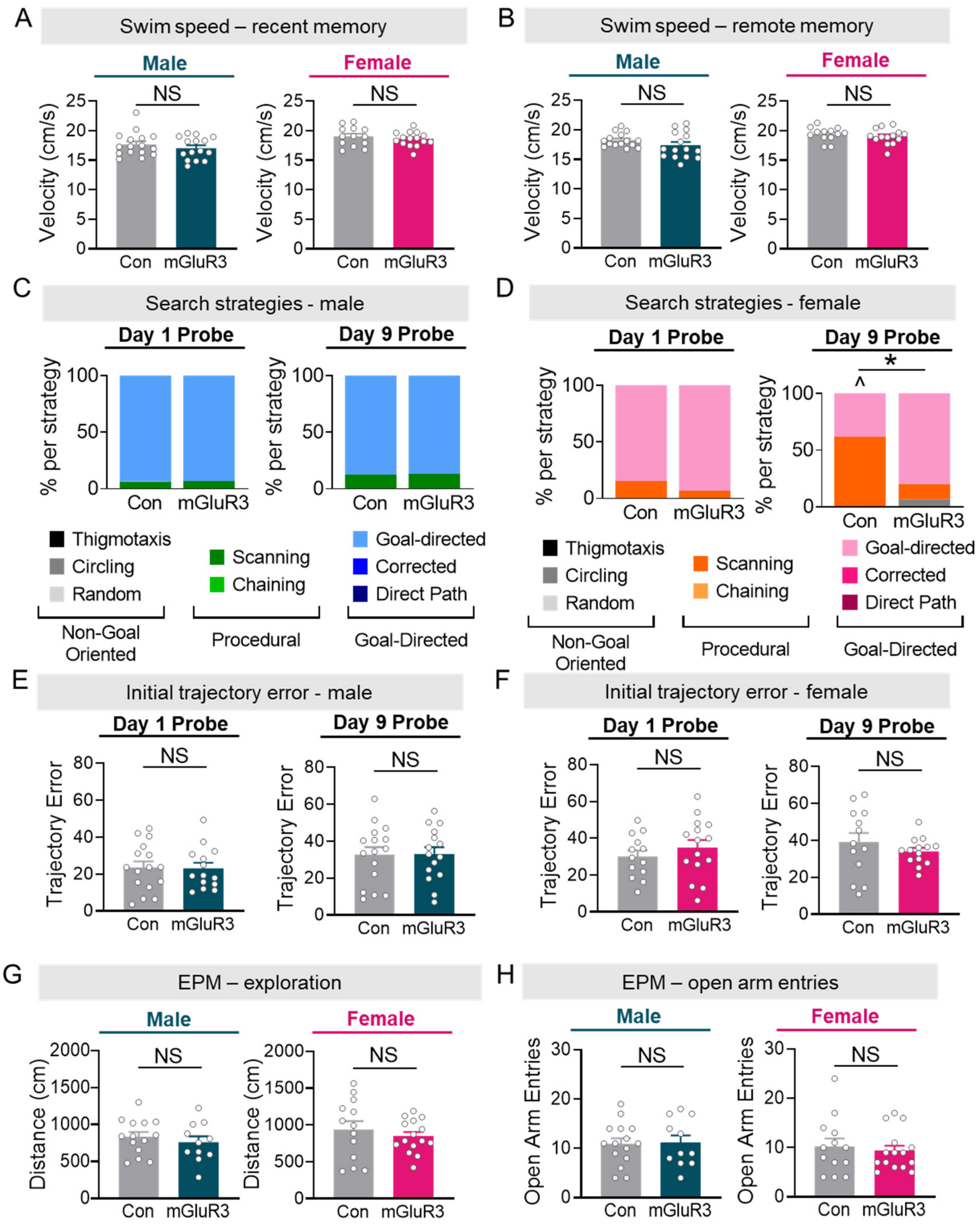


**Fig. S7. Additional behavioral characterization of *Aldh1l1*-CreERT2 mice with or without AAV vector-induced increases in astrocytic mGluR3 expression.** (**A** and **B**) Swim speeds in male and female mice during the MWM probe trial at one day (A, recent) and nine days (B, remote) post-training. (**C** and **D**) Search strategies of male (C) and female (D) mice during recent or remote probe trials quantified with Rtrack. (**E** and **F**) Initial trajectory error in male (E) and female (F) mice during recent or remote probe trials quantified with Rtrack. Data in (A–B and E–H) were analyzed with two-way ANOVA and Sidak's post-hoc test. NS = not significant. Fisher's exact test (C–D, Con vs mGluR3): *$p < 0.05$. Fisher's exact test (C and D, Female Con vs Male Con): ^$p < 0.05$.

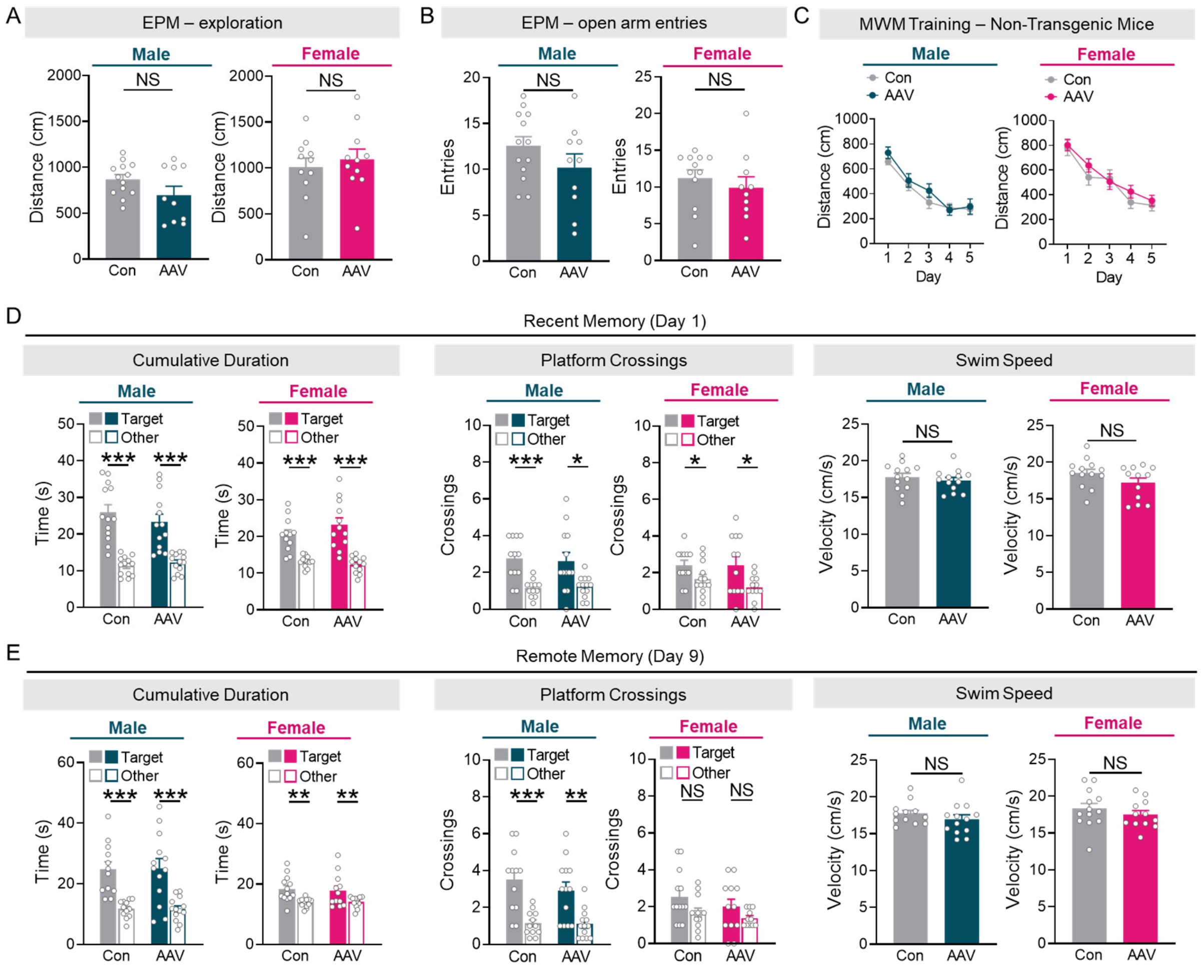


**Fig. S8. Effects of AAV vector injections on behavior in non-transgenic control littermates of *Aldh1l1*-CreERT2 mice.** (**A**) Distance traveled in the EPM by non-transgenic male and female mice with or without (Con) injection with mGluR3-encoding AAV vector. Two-way ANOVA: $F(1, 41) = 2.086$, $p = 0.1563$ for interaction effect. (**B**) EPM open-arm entries in non-transgenic male and female mice with or without AAV vector injections. Two-way ANOVA: $F(1, 41) = 0.1763$, $p = 0.6767$ for interaction effect. (**C**) Distance to reach the platform during Morris water maze training in non-transgenic mice injected with saline or Cre-dependent mGluR3-encoding AAV vector. Three-way ANOVA: $F(9, 432) = 0.7176$, $p = 0.6929$ for interaction effect. (**D**) Cumulative duration in target and non-target quadrants (two-way ANOVA: $F(1, 46) = 1.856$, $p = 0.1797$ for interaction effect); number of target and non-target platform crossings (two-way ANOVA: $F(1, 45) = 0.0286$, $p = 0.8664$ for interaction effect); swim speeds during a probe trial conducted one day after training. (**E**) Cumulative duration in target and non-target quadrants (two-way ANOVA: $F(1, 46) = 0.0431$, $p = 0.8363$ for interaction effect); number of target and non-target platform crossings (two-way ANOVA: $F(1, 46) = 0.0019$, $p = 0.9653$ for interaction effect); swim speeds during a probe trial conducted nine days after training. Data were analyzed with a two-way ANOVA (A–B and D–E) and Student's t-test (D–E). Data in (C) were analyzed with a three-way ANOVA. $^*p < 0.05$, $^{**}p < 0.01$, $^{***}p < 0.001$, NS = not significant.

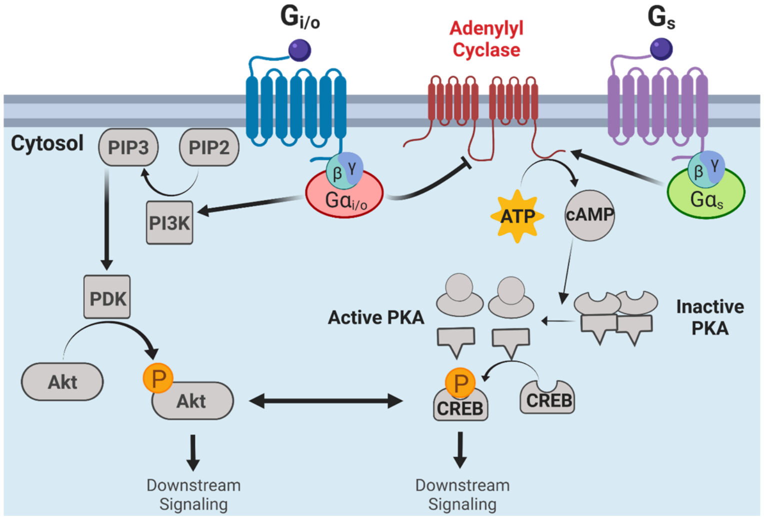


**Fig. S9.** Simplified diagram of several well-known signaling cascades engaged by $G_{i/o}$-coupled and $G_s$-coupled GPCRs and their mutual regulation.

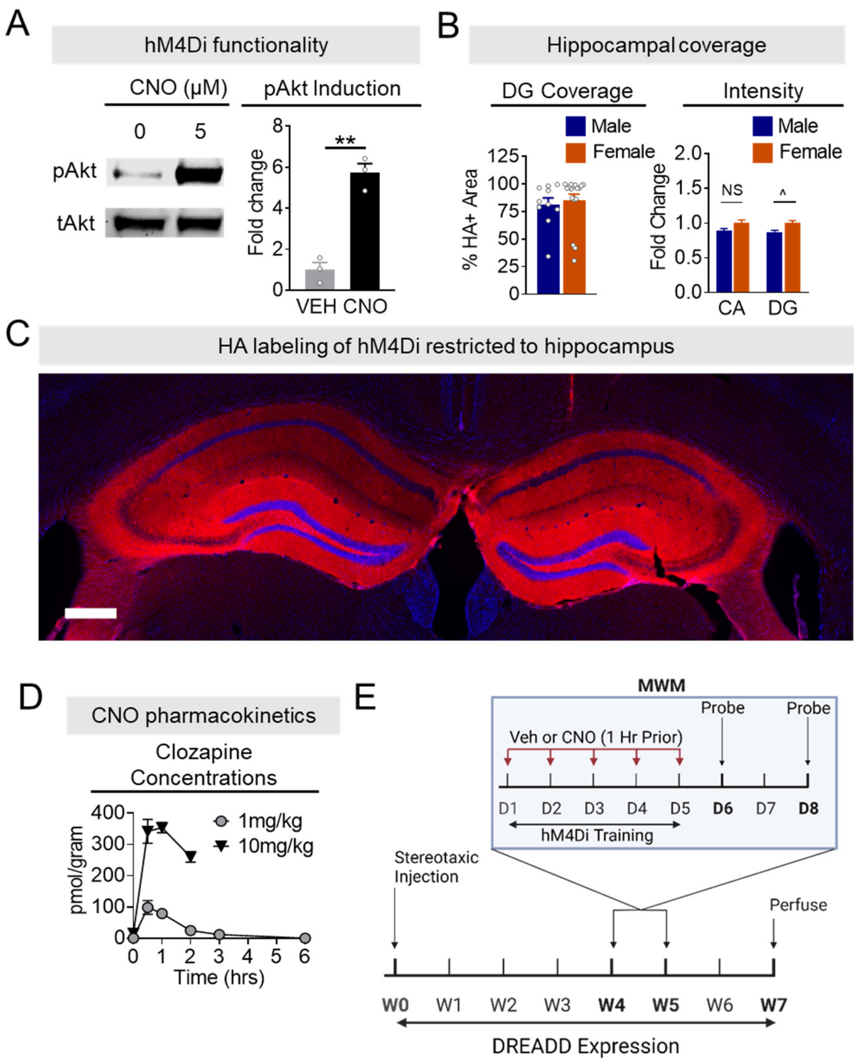


**Fig. S10. Further characterization of DREADD-encoding AAV vectors and CNO levels in the brain.** (**A**) Phospho-Akt levels following hM4Di stimulation with CNO in cultured astrocytes. Data collected from one culture. (**B**) Quantification of HA-immunopositive area in the DG and mean intensity of HA immunolabeling in the CA1 and DG regions of the hippocampus in male and female mice injected with hM4Di-encoding AAV vectors. Two-way ANOVA: $F(1, 423) = 0.0648$, $p = 0.7992$ for interaction effect. (**C**) HA-hM4Di (red) immunolabeling in hippocampal sections from *Aldh1l1*-Cre mice. HA labeling was robust in the hippocampal formation, the region where the AAVs were injected, but minimal in brain regions outside of the hippocampus. Scale bar: 400 μm. (**D**) Pharmacokinetic analyses of clozapine and CNO in the hippocampus following peripheral CNO injection (i.p.) in adult mice. In concurrence with previous findings, CNO was not detectable in the brain (data not shown), likely due to rapid conversion to clozapine (*89, 90*). (**E**) Experimental timeline for hM4Di. Data in A and B (DG coverage) were analyzed with Student's t-test: $^{**}p < 0.01$. Data in B (intensity) were analyzed with a two-way ANOVA: $^{\wedge}p < 0.05$.

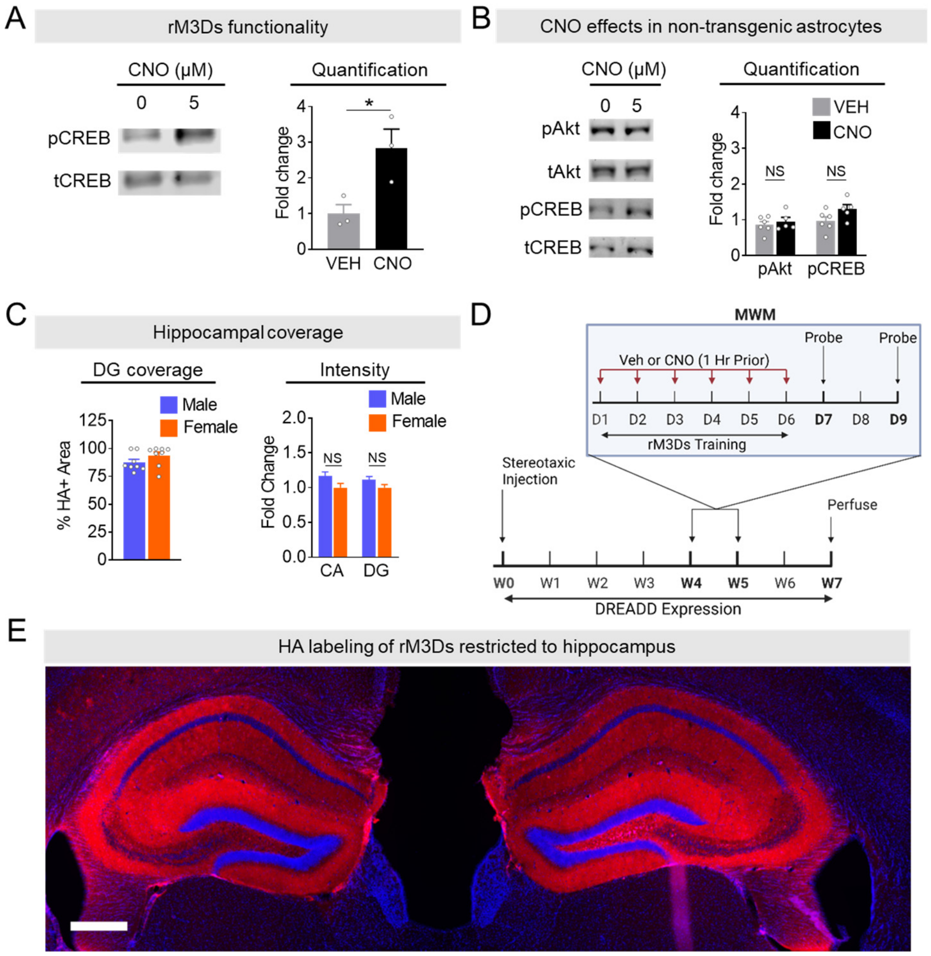


**Fig. S11. Further characterization of rM3Ds-encoding AAV vectors.** (**A**) Phospho-CREB levels following rM3Ds stimulation in cultured astrocytes. Data collected from one culture. (**B**) CNO does not cause significant increases in Akt or CREB phosphorylation levels in control astrocytes without AAV transduction. Data collected from two independent cultures. (**C**) Quantification of HA-immunopositive area in the DG and mean intensity of HA immunolabeling in the CA and DG regions of the hippocampus in male and female mice injected with rM3Ds-encoding AAV vectors. Two-way ANOVA: $F(1, 423) = 0.0648$, $p = 0.7992$ for interaction effect. (**D**) Experimental timeline for rM3Ds experiment. (**E**) HA-rM3Ds immunolabeling in hippocampal sections from *Aldh1l1*-Cre mice. HA labeling was robust in the hippocampal formation, the region where the AAVs were injected, but minimal in brain regions outside of the hippocampus. Scale bar: 400 µm. Data in A and C (DG coverage) were analyzed with Student's t-test: $*p < 0.05$. Data in B and C (intensity) were analyzed with a two-way ANOVA: NS = not significant.

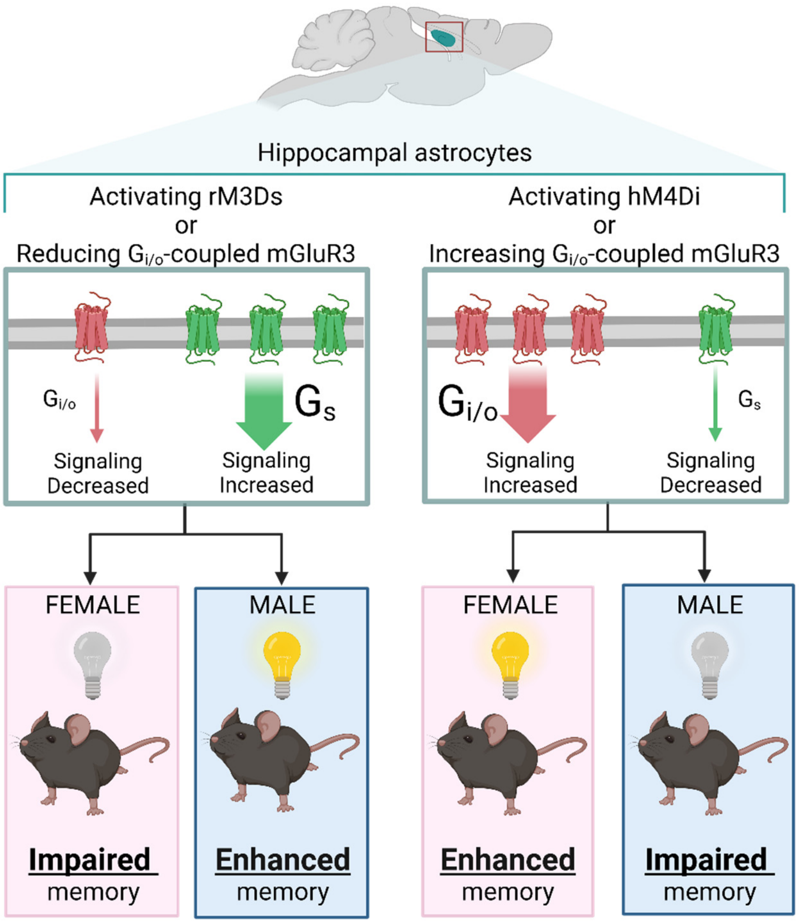


**Fig. S12. Summary of the main findings**. Alterations in astrocytic $G_{i/o}$-coupled and $G_s$-coupled receptors had bidirectional and divergent sex-specific effects on memory.

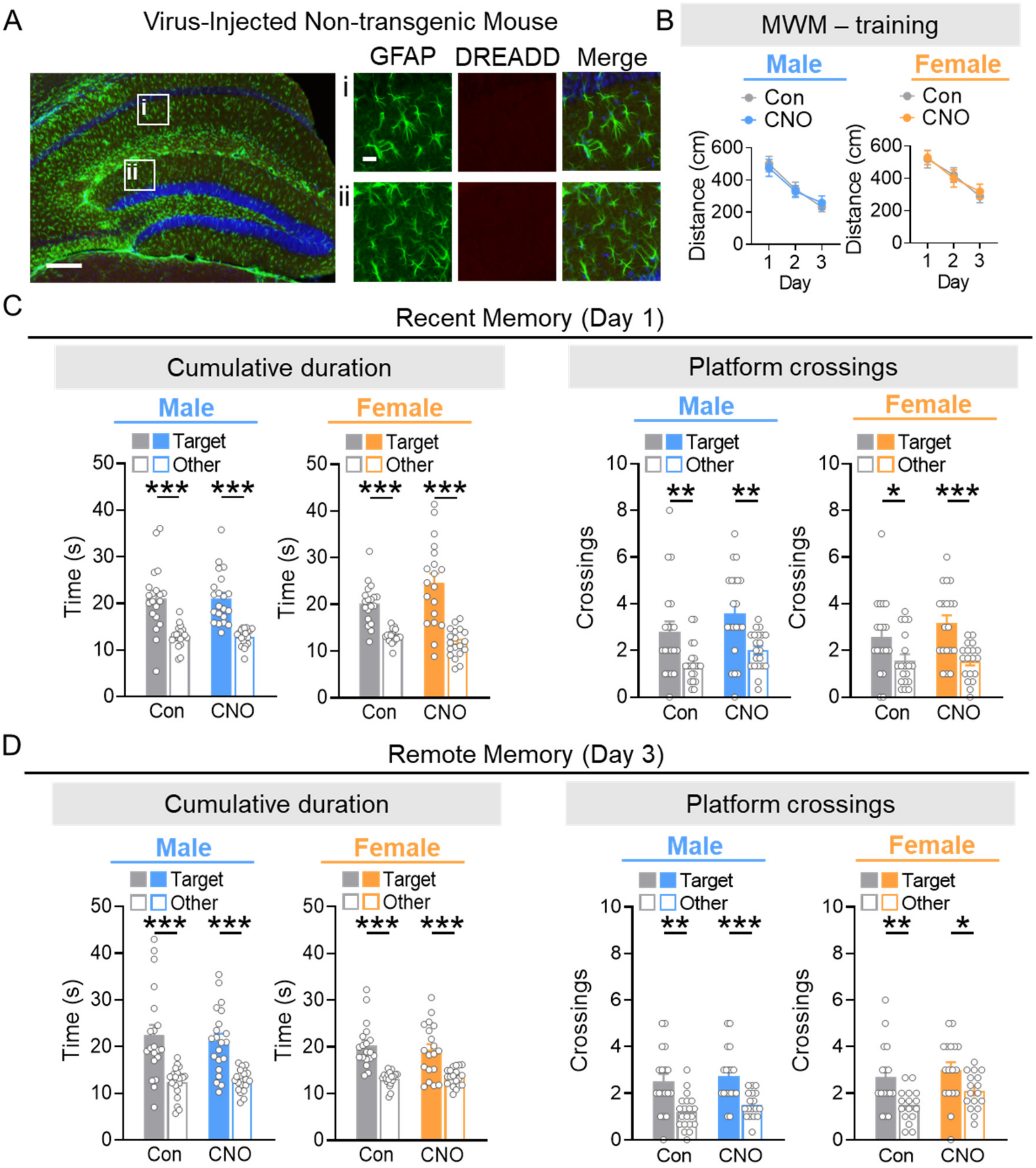


**Fig. S13. Further characterization of CNO effects in non-transgenic control mice**. (**A**) Representative images of GFAP (green) and DREADD (red) co-immunolabeling in AAV vector-injected non-transgenic mice used for testing main effects of CNO. No DREADD expression was detected in nontransgenic mice. Scale bars: 400 µm, 40 µm (inset). (**B**) Distance to reach the platform during training of non-transgenic mice injected with vehicle or CNO 1 h prior to training. Three-way ANOVA: $F(2, 111) = 0.07668$, $p = 0.9262$ for interaction effect (time x sex x CNO). (**C**) Target quadrant preference and platform crossings compared to other analogous locations during a probe trial performed one day after training. Mixed-effects ANOVA: $F(1, 37) = 2.714$, $p = 0.1079$ for interaction effect (target duration). Mixed-effects: $F(1, 37) = 0.0614$, $p = 0.8057$ for interaction effect (target crossings). (**D**) Target quadrant preference and platform crossings compared to other analogous locations during a probe trial performed three days after training. Mixed-effects: $F(1, 37) = 0.0038$, $p = 0.9513$ for interaction effect (target duration). Mixed-effects: $F(1, 66) = 0.2836$, $p = 0.5961$ for interaction effect (target crossings) Data in (B) were analyzed with a three-way ANOVA. Data in (C and D) were analyzed with a two-way ANOVA and Student’s t-test. *$p < 0.05$, **$p < 0.01$, ***$p < 0.001$.

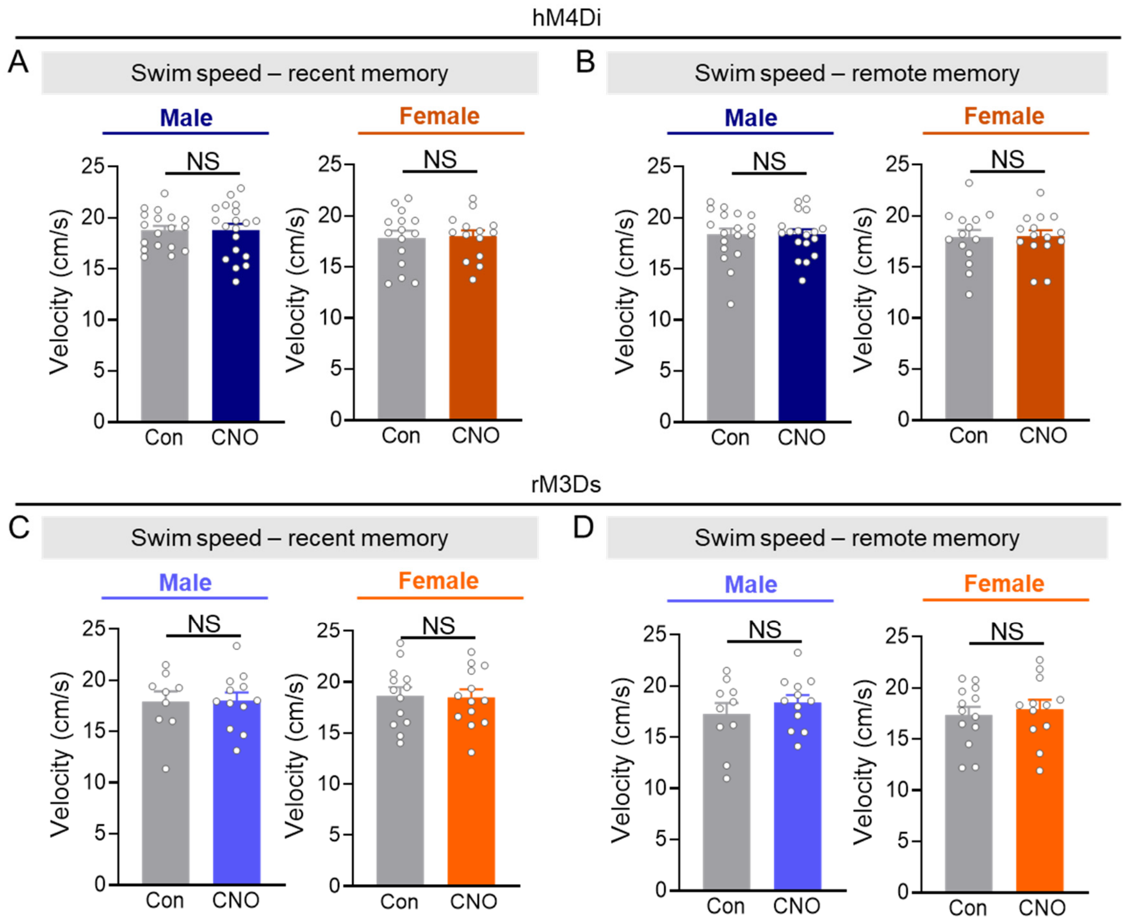


**Fig. S14. Swim speeds in mice with chemogenetic manipulations.** Swim speeds in Morris water maze probe trials conducted one (**A** and **C**) or three (**B** and **D**) days post-training. (**A** and **B**) Swim speeds in mice expressing astrocytic hM4Di in hippocampal astrocytes. Two-way ANOVA: $F(1, 62) = 0.0128$, $p = 0.9105$ for interaction effect (recent), $F(1, 61) = 0.0070$, $p = 0.9334$ for interaction effect (remote). (**C** and **D**) Swim speeds in mice expressing astrocytic rM3Ds in hippocampal astrocytes. Two-way ANOVA: $F(1, 43) = 0.0282$, $p = 0.8674$ for interaction effect (recent), $F(1, 43) = 0.0953$, $p = 0.7590$ for interaction effect (remote). All data were analyzed with a two-way ANOVA and Sidak's post-hoc test. NS = not significant.

| Figure # | Panel | Condition | Sex | |
|---|---|---|---|---|
| | | | *Male* | *Female* |
| 1 | C (CA) | Con | 6 | 6 |
| | | sgRNA | 4 | 5 |
| | C (DG) | Con | 6 | 6 |
| | | sgRNA | 4 | 5 |
| | D (CA) | Con | 7 | 11 |
| | | sgRNA | 8 | 7 |
| | D (DG) | Con | 8 | 9 |
| | | sgRNA | 5 | 10 |
| | F-H | Con | 12 | 12 |
| | | sgRNA | 10 | 11 |
| 2 | C | Con | 12 | 11 |
| | | mGluR3 | 13 | 15 |
| | D | Con | 12 | 10 |
| | | mGluR3 | 13 | 14 |
| | E-G | Con | 16 | 13 |
| | | mGluR3 | 15 | 15 |
| 3 | D-F | Con | 19 | 15 |
| | | CNO | 19 | 15 |
| 4 | D-F | Con | 12 | 13 |
| | | CNO | 12 | 13 |

**Table S1.** Numbers of mice used in each experiment shown in the main figures.

| Figure # | Panel | Condition | Sex | |
|---|---|---|---|---|
| | | | *Male* | *Female* |
| S1 | D | | 14 | 12 |
| | E (CA1 + DG) | GFAP | 15 | 12 |
| | | Iba1 | 17 | 15 |
| | | NeuN | 6 | 6 |
| S2 | B | Con | 11 | 9 |
| | | sgRNA | 7 | 8 |
| | C (protein) | | 6 | 6 |
| | C (RNA) | | 9 | 11 |
| | D | Con | 10 | 9 |
| | | sgRNA | 8 | 10 |
| S3 | A-F | Con | 12 | 12 |
| | | sgRNA | 10 | 11 |
| | G | Con | 11 | 11 |
| | | sgRNA | 9 | 10 |
| | H | Con | 12 | 12 |
| | | sgRNA | 10 | 10 |
| S6 | A | | 9 | 9 |
| | D | | 6 | 5 |
| S7 | A-F | Con | 16 | 12 |
| | | mGluR3 | 15 | 14 |
| | G-H | Con | 16 | 13 |
| | | mGluR3 | 14 | 15 |
| S8 | A-E | Con | 13 | 13 |
| | | mGluR3 | 14 | 12 |
| S10 | B | | 11 | 20 |
| | D | 1mg/kg | 3-4/timepoint | |
| | | 10mg/kg | 3-4/timepoint | |
| S11 | C | | 10 | 10 |
| S13 | B-D | Con | 21 | 19 |
| | | CNO | 21 | 19 |
| S14 | A-B | Con | 19 | 15 |
| | | CNO | 19 | 15 |
| | C-D | Con | 12 | 13 |
| | | CNO | 12 | 13 |

**Table S2.** Numbers of mice used in each experiment shown in the supplemental figures.

| Gene | Forward Primer | Reverse Primer |
|---|---|---|
| Grm1 | GCAGCGAGCCTTGCTTAAA | AGATCCAGCAGCAGCTCAC |
| Grm2 | GGACTTCGTGCTCAATGTCA | CCATCTCCAAAGCGGTCAAA |
| Grm3 | ACCTCAACAGGTTCAGTGTCA | TTGCACACTGTCGGGACATA |
| Grm4 | TCAAGAAGGGAAGCCACATCAA | ACCTTCCCCTCCTGTTCGTA |
| Grm5 | TGCAGTGAACCGTGTGAGAA | AAGGTGTGCAGGTCCAACAA |
| Grm7 | AAGGAGCCATCACCATCCAA | TCAAGTGTCCGGGATGTGAA |
| Grm8 | CCACTGGACCAATCAACTTCAC | GGGTGCGTGTGCTCTCTATTA |
| Actb | CCCTAAGGCCAACCGTGAAA | AGCCTGGATGGCTACGTACA |
| Gapdh | CAAGGTCATCCCAGAGCTGAA | CAGATCCACGACGGACACA |
| Gusb | AGTATGGAGCAGACGCAATCC | ACAGCCTTCTGGTACTCCTCA |
| Tbp | CCTTGTACCCTTCACCAATGAC | ACAGCCAAGATTCACGGTAGA |

**Table S3.** Mouse gene primer sequences used for RT-qPCR.
